# From Energy to Cellular Force in the Cellular Potts Model

**DOI:** 10.1101/601781

**Authors:** Elisabeth G. Rens, Leah Edelstein-Keshet

## Abstract

Single and collective cell dynamics, cell shape changes, and cell migration can be conveniently represented by the Cellular Potts Model, a computational platform based on minimization of a Hamiltonian while permitting stochastic fluctuations. Using the fact that a force field is easily derived from a scalar energy (**F** = −∇*H*), we develop a simple algorithm to associate effective forces with cell shapes in the CPM. We display the predicted forces for single cells of various shapes and sizes (relative to cell rest-area and cell rest-perimeter). While CPM forces are specified directly from the Hamiltonian on the cell perimeter, we infer internal forces using interpolation, and refine the results with smoothing. Predicted forces compare favorably with experimentally measured cellular traction forces. We show that a CPM model with internal signaling (such as Rho-GTPase-related contractility) can be associated with retraction-protrusion forces that accompany cell shape changes and migration. We adapt the computations to multicellular systems, showing, for example, the forces that a pair of swirling cells exert on one another, demonstrating that our algorithm works equally well for interacting cells. Finally, we show forces associated with the dynamics of classic cell-sorting experiments in larger clusters of model cells.

**Author summary:** Cells exert forces on their surroundings and on one another. In simulations of cell shape using the Cellular Potts Model (CPM), the dynamics of deforming cell shapes is traditionally represented by an energy-minimization method. We use this CPM energy, the Hamiltonian, to derive and visualize the corresponding forces exerted by the cells. We use the fact that force is the negative gradient of energy to assign forces to the CPM cell edges, and then extend the results to interior forces by interpolation. We show that this method works for single as well as multiple interacting model cells, both static and motile. Finally, we show favorable comparison between predicted forces and real forces measured experimentally.

## Introduction

From embryogenesis and throughout life, cells exert forces on one another and on their surroundings. Cell and tissue forces drive cell shape changes and cell migration by regulating cell signaling and inducing remodeling of the cytoskeleton. Along with progress in experimental quantification of cellular forces, there has been much activity in modeling and developing computational platforms to explore cellular mechanobiology. In some platforms, among them vertex-based and cell-center based simulations, the shape of a cell is depicted by convex polygons, ellipsoids or spheres.

The Cellular Potts Model (CPM) is a convenient computational platform that allows for a variety of irregular and highly fluctuating cell shapes. Unlike vertex-based computations, the CPM easily accommodates cell detachment or reattachment from an aggregate, and a range of cell-cell adhesion. It also captures stochastic aspects of cell movement. At the same time, since CPM computations are based on a phenomenological “energy”, the Hamiltonian, it has often been criticized as non-physical, or, at least, as devoid of Newtonian forces. In their review of models for cell migration, Sun and Zaman [1] point to the need to coordinate results between force-based and energy-based models, indicating that this is a “challenging but significant” problem. Here we devise a map between the CPM Hamiltonian and an explicit vector-field of forces associated with the dynamics of cell shape. We illustrate the computation of this force field for single cells, for pairs of cells, and for larger clusters of cells interacting through adhesion and through internal signaling.

In the Cellular Potts model, each “cell” configuration *σ*, consists of *N* lattice site, assigned a unique index (“spin number”). Parts of the domain containing no cells are indexed 0 by convention. For a single CPM cell surrounded by medium, the typical Hamiltonian is given by

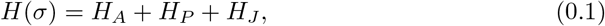

where σ is the cell configuration and

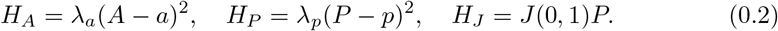

Here *H_A_* is an energetic cost for expansion or contraction of the area, *A*, away from a constant “rest area”, *a*, of the cell. *H_P_* is energetic cost for deviation of the cell perimeter *P* from its “rest perimeter” *p. H_J_* is energy associated with the cell-medium interface (generalized later to include cell-cell or cell-medium adhesive energies.) The factors λ_*a*_, λ_*p*_, *J*(0,1) set the relative energetic costs of area changes, perimeter changes, and changes in the contact with the medium. In a typical CPM simulation, cell shapes are highly deformable. At each simulation step (Monte Carlo Step, MCS) every boundary pixel of each cell may “protrude” or “retract”. Formally, these changes are denoted “spin-flips”, and are accepted or rejected with some probability that depends on the change in *H* and on a user-defined “temperature” *T*, as described in Materials and Methods.

There are many realizations of the Potts Model with additional terms, or variations of such terms. In the Discussion, we summarize the numerous ways that CPM cell shape computations were linked to force calculations external to the CPM formalism itself (and including, among others, finite element methods).

Since the Hamiltonian associates an “energy” with each cellular configuration, theoretically we can relate forces to the negative gradient of the Hamiltonian, i.e.

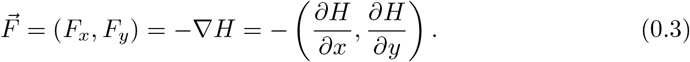

In practice, the computations are all carried out on a finite grid, so partial derivatives in (0.3) are approximated by finite differences. We do this by calculating the small change in the Hamiltonian when the cell boundary is extended in the *x* or *y* directions by a small step Δ*x* or Δ*y*, as illustrated in Fig. 1. This is repeated at each point along the edge of the cell.

**Figure 1.**
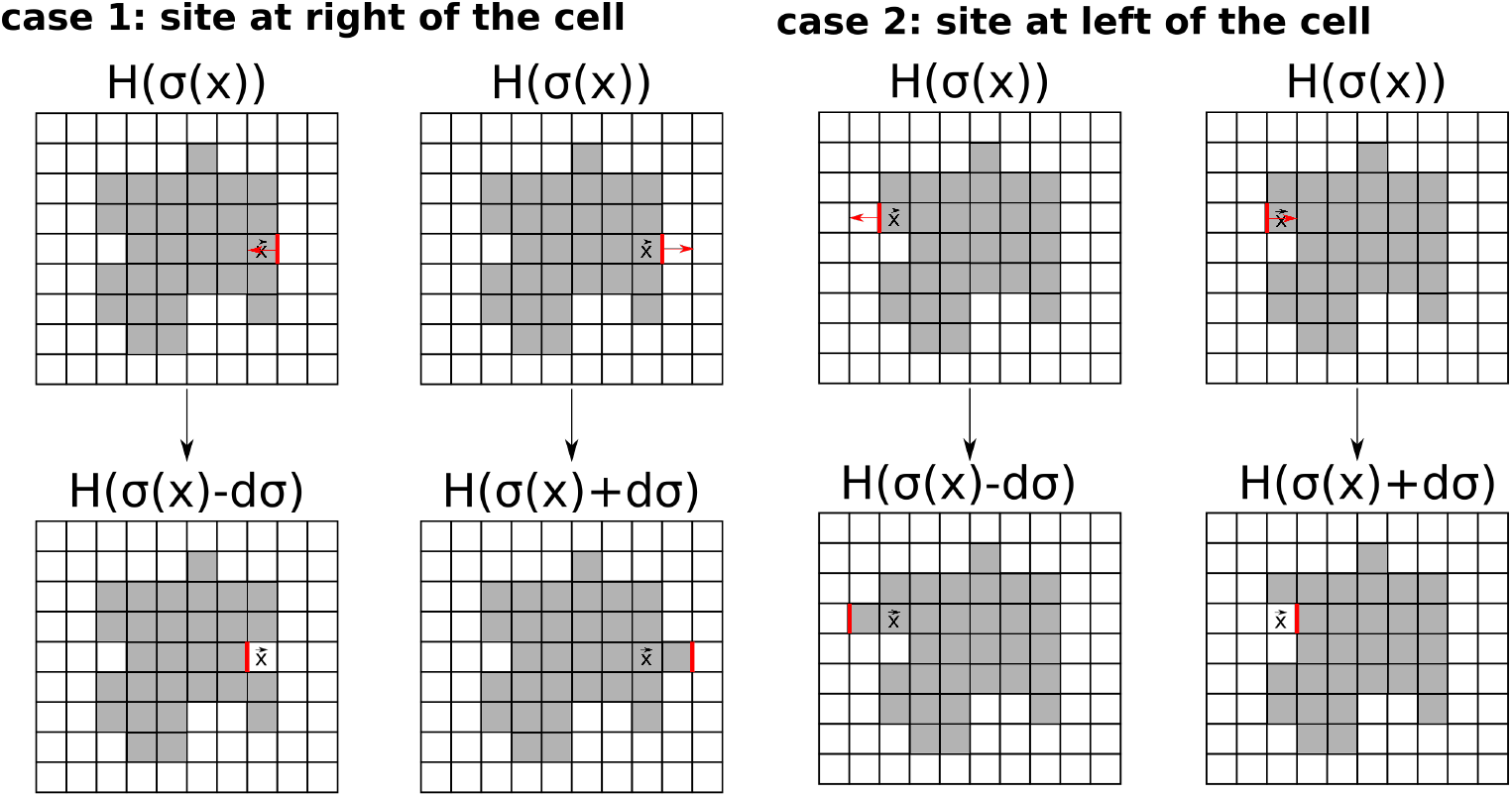
Schematic diagram showing how forces are derived from a Cellular Potts Model Hamiltonian. The Hamiltonian represents an energy, so 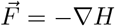. We compute a discrete approximation to the components of the force (*F_x_, F_y_*) at each point 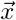 on the cell boundary. Centered finite differences are used to approximate the derivatives –(*∂H/∂x,∂H/∂y*) of the Hamiltonian as in Eq. (0.5). Here we illustrate the idea for the *x* component of the force, *F_x_*. From a given initial CPM cell configuration *σ* (top row), we numerically compute the difference in the Hamiltonian at a point 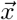 on the right cell boundary when the cell retracts (left most column) or extends (next column). We show the same idea for the left cell boundary (next two columns). The force field computed along the boundary is then smoothed and interpolated to the cell interior, as described in Materials and Methods.

A feature of CPM is that it only associates energetic costs with small fluctuations of the edge of a model cell. Hence, such computation results in forces along the cell perimeter that must then be interpolated elsewhere. The workflow then entails 1) calculating the force along the cell perimeter, 2) reducing the grid effect in the force field, 3) interpolating the force-field to the interior of the cell. This generic computation can be extended to forces of multiple interacting cells (in a cell sheet or aggregate). The implementation of this idea is described in the Materials and Methods Section, with further details in the Supplementary Information.

## Results

### Forces associated with static cell shapes

We computed the force-fields associated with the CPM Hamiltonians of single static cells with circular (A), elliptical (B), and irregular shapes (C,D). Results of the complete algorithm (including smoothing and interior forces) are shown in Figure 2 (A-D). Intermediate calculations (forces on the cell boundary without and with smoothing, and without interior smoothing) can be found in the Supplementary Figures 14–16.

**Figure 2.**
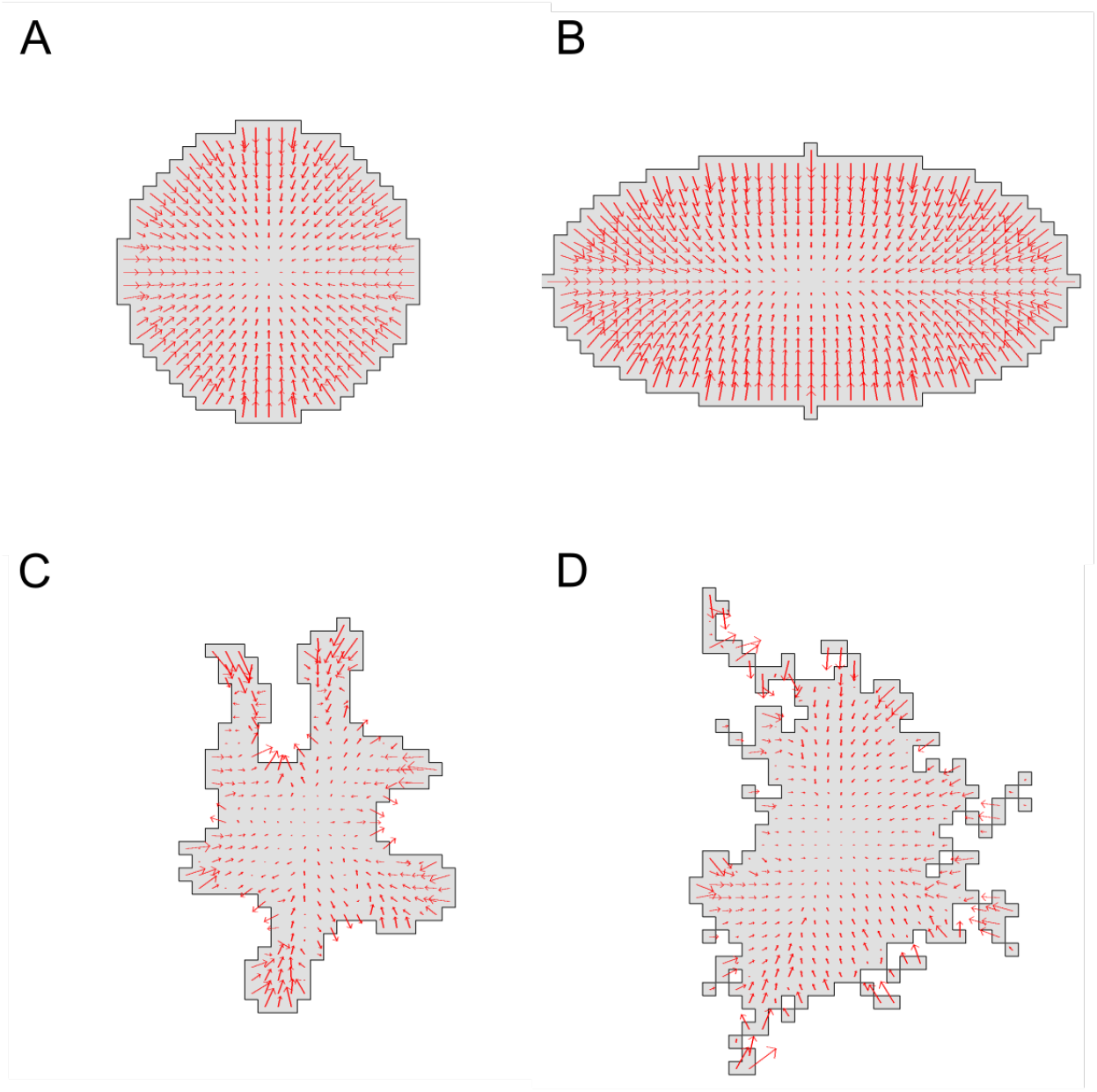
Forces predicted for several cell shapes. Force fields predicted by our complete method (smoothing and internal interpolation) for four simulated cell shapes in the CPM. (A) Circular cell (area *A* = 401, perimeter *P* = 74, diameter=23). (B) Elliptical cell (area *A* = 629, perimeter *P* = 101, axes lengths 21 and 41). (C) Irregular cell shape (area *A* = 301, perimeter *P* = 118). (D) Highly irregular cell shape (area *A* = 400, perimeter *P* = 146). Parameter values were *a* = 300, λ_*a*_ = 10, *p* = 100, λ_*p*_ = 10, *J*(0,1) = 3000, *ξ*(*r*) = 18, and *r* = 3 for all neighborhood calculations. We used a grid of 50 by 50 lattice sites with Δ*x*=1. See also Supplementary Figures 14–16 for intermediate steps in the calculation of forces.

Whether forces point inwards or outwards depends on the values of the area A and perimeter *P* relative to their target values *a,p* and the relative weights of the energetic cost or area and perimeter changes. For parameters given in Fig. 2, forces point inwards all along the boundary of the circular and elliptical cell shapes. We find forces directed approximately normal to the boundary, with magnitudes that decay towards the centroid, as a consequence of our interpolation.

In more irregular shapes (Figure 2C,D), forces can point either inwards or outwards at different points along the boundary. For the irregular cell with given configuration and Hamiltonian parameters, we found that at convex sites, the forces point inwards, while at concave sites, the forces point outwards. This is also in line with expectations based on local (positive or negative) curvature of the boundary. Even for the most irregular cell shape, the forces are fairly smooth and continuous.

### Dynamic cell shapes and evolving forces

We next tracked the evolution of forces that accompany dynamic changes in shape of a CPM “cell”. To do so, we initiated a CPM computation with a circular cell with perimeter smaller than the rest-length *p* and area greater than the rest area, *a*. We also assumed λ_*p*_ > λ_*a*_, so that the energetic cost of the perimeter term dominated the energetic cost of the area term in the Hamiltonian.

A time sequence of cell shapes and accompanying forces is shown in Fig. 3. At MCS step 1, the cell is far from its preferred configuration, and large forces are seen all along its edge. (Note that these forces are mostly directed outwards, with notable exceptions in non-convex regions of the boundary.) As the cell quickly obtains its target perimeter, the forces point inwards and the cell starts to shrink to obtain its target area. The irregular force directions and large magnitudes rapidly decline, so that by MCS 3, the force-field is more regular, and directed “inwards”. The cell becomes highly ramified, with thin protrusions so as to satisfy both area and perimeter constraints. After a few MCS, the cell shape has equilibrated, and forces decrease to very low levels.

**Figure 3.**
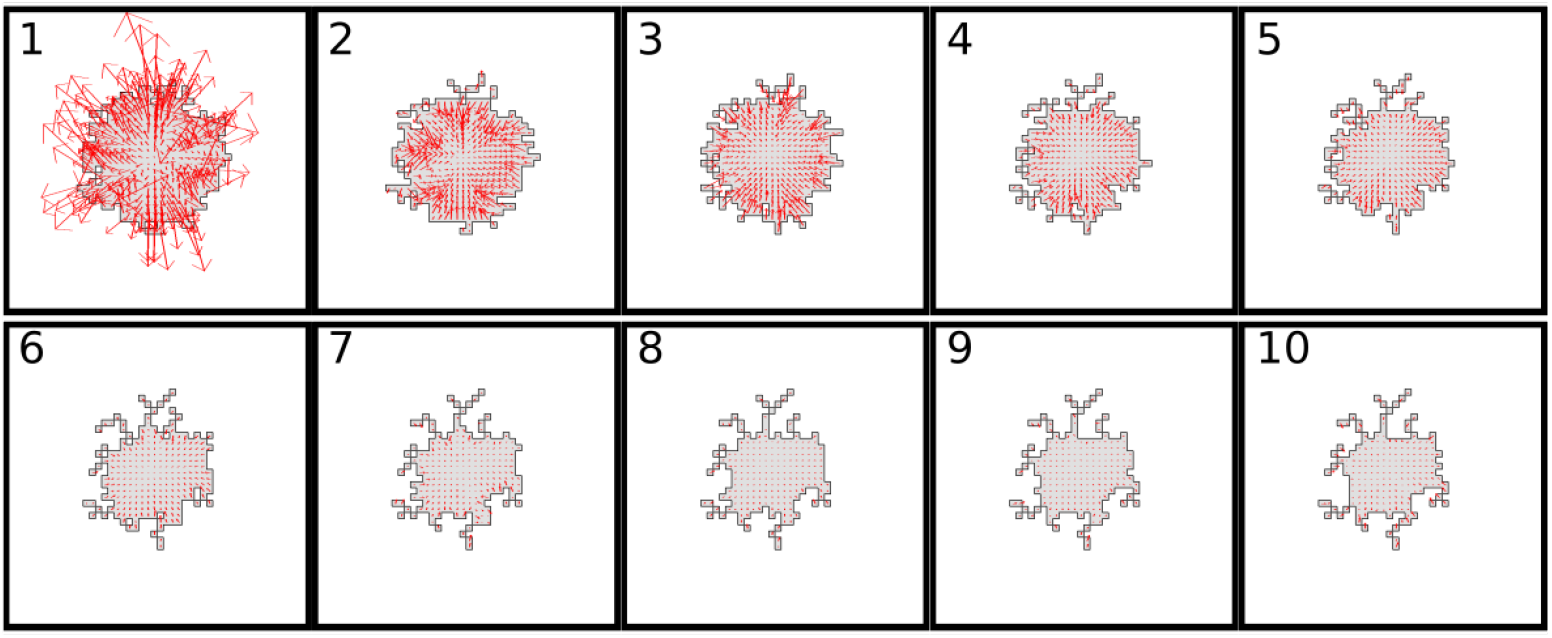
Dynamics of cell shape and the evolution of forces. Time series from 1 to 10 MCS. The cell achieves force balance by decreasing its area and increasing its perimeter. Parameters were *a* = 200, λ = 8, *p* = 100, λ_*p*_ = 2000, *J*(0,1) = 3000, *T* = 10. The cell areas *A* at each of the 10 Monte Carlo steps displayed are: *A* = 397, 364, 332, 299, 280, 250, 232, 219, 213, 208, and the perimeter is *P*=74, 94.8, 98.2, 100.1, 99, 99.2, 99.1, 98.9, 99, 99.4.

### Active forces from internal signaling

Several models have proposed signaling kinetics inside cells that result in forces of protrusion or retraction (powered by actin assembly or actomyosin contractility). Among these is the simple “wave-pinning” model for GTPase spatio-temporal distribution [6]. We sought to visualize the evolution of the internal GTPase activity field in parallel with the forces that it creates. Consequently, we assumed that a single signaling protein in two states (analogous to active and inactive RhoA) participates in reaction-diffusion kinetics inside the deforming “cell” and leads to edge contraction.

Regions of high Rho activity contiguous to the cell edge are shaded light green in Fig. 4A. The internal chemistry leads to a force of protrusion, modeled by an additional term, Δ*H_p_*, superimposed on the Hamiltonian change.We assume Δ*H_p_* = ±*βu*, where *β* > 0 is a constant, and *u* the local signaling activity level. We assume that the signal promotes contraction, so that Δ*H_p_* is negative for retractions and positive for protrusion.

**Figure 4.**
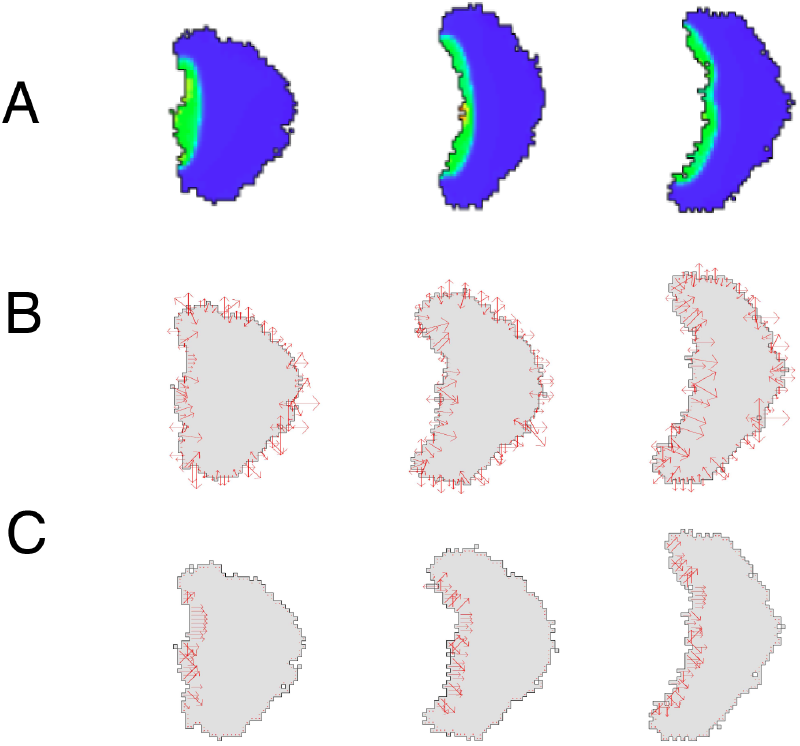
Active contractile forces from internal signaling. Shown is a time sequence (left to right, at 50, 100 and 150 MCS) of a moving cell whose shape changes in response to a polarizing internal signal (e.g. Rho GTPase). (A) The internal GTPase field (bright green at high values) based on the Wave-Pinning model. High levels of activity are assumed to create large local inwards contraction. (B) Total forces given by ∇*H* + *dH_G_* along the perimeter of the deforming cell. (C) Forces due to the active contraction term (*dH_G_*). Forces are shown without smoothing or interpolation. Parameter values for CPM were: λ_*a*_ = 10, *a* = 1000, λ_*p*_ = 0, *J*(0,1) = 5000, *T* = 50; parameter values for internal signaling: *β* = 40 (for *dH_p_*), the numerical redistribution radius was *r* = 3 (active rho), *r* = 75 (inactive rho). Parameters for internal reaction-diffusion system, and details for the numerical method are provided in the Supplementary Information.

Results are shown in Fig. 4A-C as a time sequence of cell deformations from left to right. In Fig. 4A, we see that chemical polarization is maintained, as described in previous studies [7,10–12]. Contraction of the cell rear leads to the expansion of other cell edges based on the CPM area constraint. In Figure 4B and C, the total force field and the protrusive forces respectively are shown. Due to high signaling levels at the left edge of the cell, a contractile force pointing towards the right develops (Figure 4C). At the right side of the cell, forces due to the area and perimeter constraints point outwards. All in all, these forces result in migration of the cell to the right.

### Comparison with experimental force fields

Single cells can apply significant forces that remodel the extracellular matrix. In traction force microscopy, beads are embedded into a substrate on which cells adhere. By tracking bead displacements, cell traction forces can be inferred. Such inverse methods quantify and reveal very detailed force fields. Traction forces roughly point towards the cell’s centroid, are highest in protruding regions and decline towards the cell’s centroid [8,13–16]. Via cell-cell adhesions, cells also apply forces on neighbouring cells and these forces can propagate through tissues [17].

We asked how the predictions of the CPM-based force fields compare with data for actual traction forces observed in real cell experiments. Consequently, we utilized data kindly provided by the authors of [8] for two cancer cell lines. Several steps were needed to arrive at a shared grid, to select CPM parameters, and to compare magnitudes on a similar range, and adjust smoothing. Details are described in Methods and in the Supplementary Information. Two examples are shown in Figs. 5–6. Similar results (not shown) were obtained with other data from the same paper.

**Figure 5.**
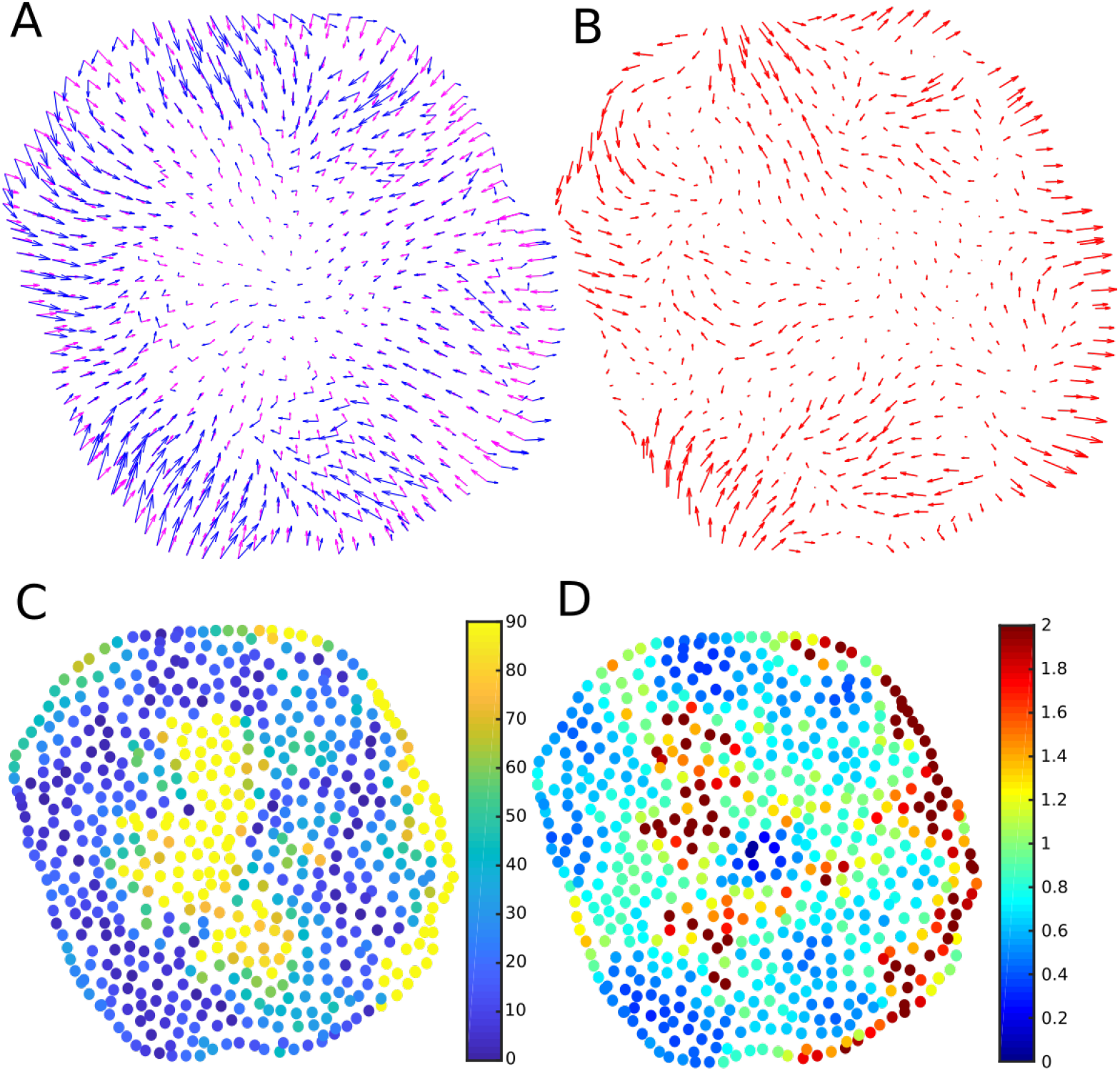
Comparing predicted forces to experimental data for a round cell. (A) Predicted CPM force fields (magenta arrows) and experimental data (blue arrows) (B) Difference of CPM force field and data force field (C) directional deviation (angle between predicted and experimentally observed force vectors), dark blue means forces align well. (D) relative magnitudes of the force fields, green means similar magnitude. Parameter values are given in Table 1.

**Figure 6.**
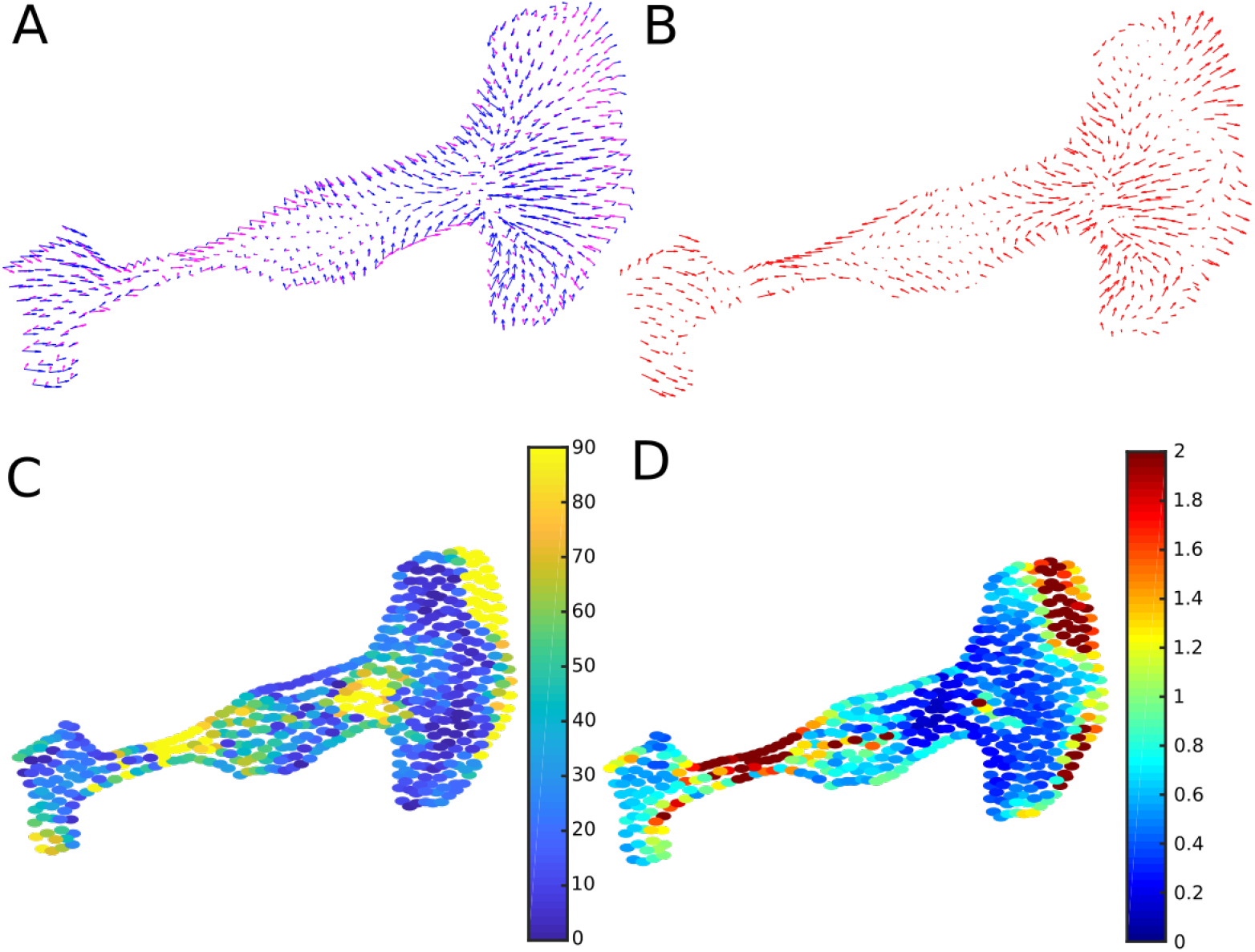
Comparing predicted forces to experimental data for a polarized cell. Panels as in Fig 5 but for a polarized cell Parameter values are given in Table 2.

Figs. 5A and 6A show observed (blue) and CPM predicted (magenta) force fields superimposed on the same grid. Overall, we find surprisingly good qualitative agreement, given the simplicity of the method. Experimental and predicted forces point roughly in the same direction, for much of the cell shape. The concordance is particularly good for the round cell, where our approximation for centroid-pointing internal forces appears to be quite good. For the polarized cell in Fig. 6, this agreement is less accurate, as two distinct “foci” appear to organize the force field in the experimental data. Figs. 5B and 6B show the difference, *F_data_* – *F_CPM_*. As expected, there are regions in each cell where localized internal forces (not captured by CPM) result in significant deviation between data and predictions.

We compared directions of predicted and experimental forces at corresponding points. Results are shown in Figure 5C and Figure 6C, with dark blue for points where observed and predicted forces are aligned, and yellow-orange for points at which the predicted direction deviates strongly from its observed value. Within a reasonable range of the cell edge, the model captures the direction of the forces very well. This correspondence is quantified in Supplementary Figure 22A. In the interior of the cell, force magnitudes are so small that directions carry large errors, and we cannot judge accuracy of the predictions.

We also compared relative force magnitudes, by plotting |*F_CPM_*|/|*F_data_*| in Figures 5D and 6D. We find some regions of deviation, notably at the top right corner of the spindle-shaped cell. Supplementary Figure 22B shows the overall comparison of force magnitude deviation.

We tested a variety of CPM parameter values, including those that provide optimal L2 norm fits of predictions to data (See Tables 1 and 2). A comparison of results for distinct CPM parameters is shown in Supplementary Figures 19 and 20. The ‘optimal’ CPM parameter values vary over a much larger range for the round cell than for the polarized cell data. There are many parameters λ, λ_*p*_, *A, P* and *J*(0,1) that determine the overall magnitude of the force, so it is not surprising that a good fit is obtainable with different values.

Finally, we also display a time series of cell movements in Figure 21 comparing the CPM force field with the data for the polarized cell. During active cell motion, large traction forces are built up for translocation (long blue arrows in the protrusive front of the cell in Figure 21 C, D. The basic CPM cannot account for these forces, which are due to active contractile or protrusive elements in the cell. However, as shown in a previous section, it would be possible to decompose a Hamiltonian into components corresponding to active forces and shape-based forces. In a similar vein, the difference of the forces *F_data_* – *F_CPM_* could provide an estimate for the spatially distributed forces of active protrusion/contraction in a motile cell.

### Interacting cells and adhesion forces

Forces between interacting cells are not easy to measure directly. However, they have been inferred from high-resolution traction-force measurements, for example by [18] using a force-balance principle and thin-plate FEM analysis.

Here, we asked whether our algorithm would predict intercellular forces in two or more cell that interact by adhesion. To investigate this question, we considered two scenarios, including simple adhesion and signaling-regulated motility in a pair of cells. Results are given below. Note that in the CPM, a high adhesive energy cost J, corresponds to low cell-cell adhesion.

### Varying adhesion strength

For the adhesion experiment, we set λ_*p*_ = 0, to omit the perimeter constraint, and used only the target area and adhesive energy in the Hamiltonian. We explored several values of the adhesive energy *J*(1, 2) between cells, keeping both cells equally adherent to the ‘medium’, *J*(0,1) = *J*(0, 2) = constant. Results are shown in Figure 7. We find that for highly ‘sticky’ cells (*J*(1, 2) < 2*J*(0,1)), the cells remain attached with a wide contact region, as shown in Fig. 7A. In the neutral cell-cell adhesion case, *J*(1, 2) = 2*J*(0, 1), in Figure 7B, the cells remain attached on a smaller contact interface. In this case, the round green cell initially (at 1 MCS), applies the same force magnitudes at every interface (note circled regions). Finally, in 7C, with *J*(1, 2) > 2*J*(0,1), the energetically favored configuration is detached cells.

**Figure 7.**
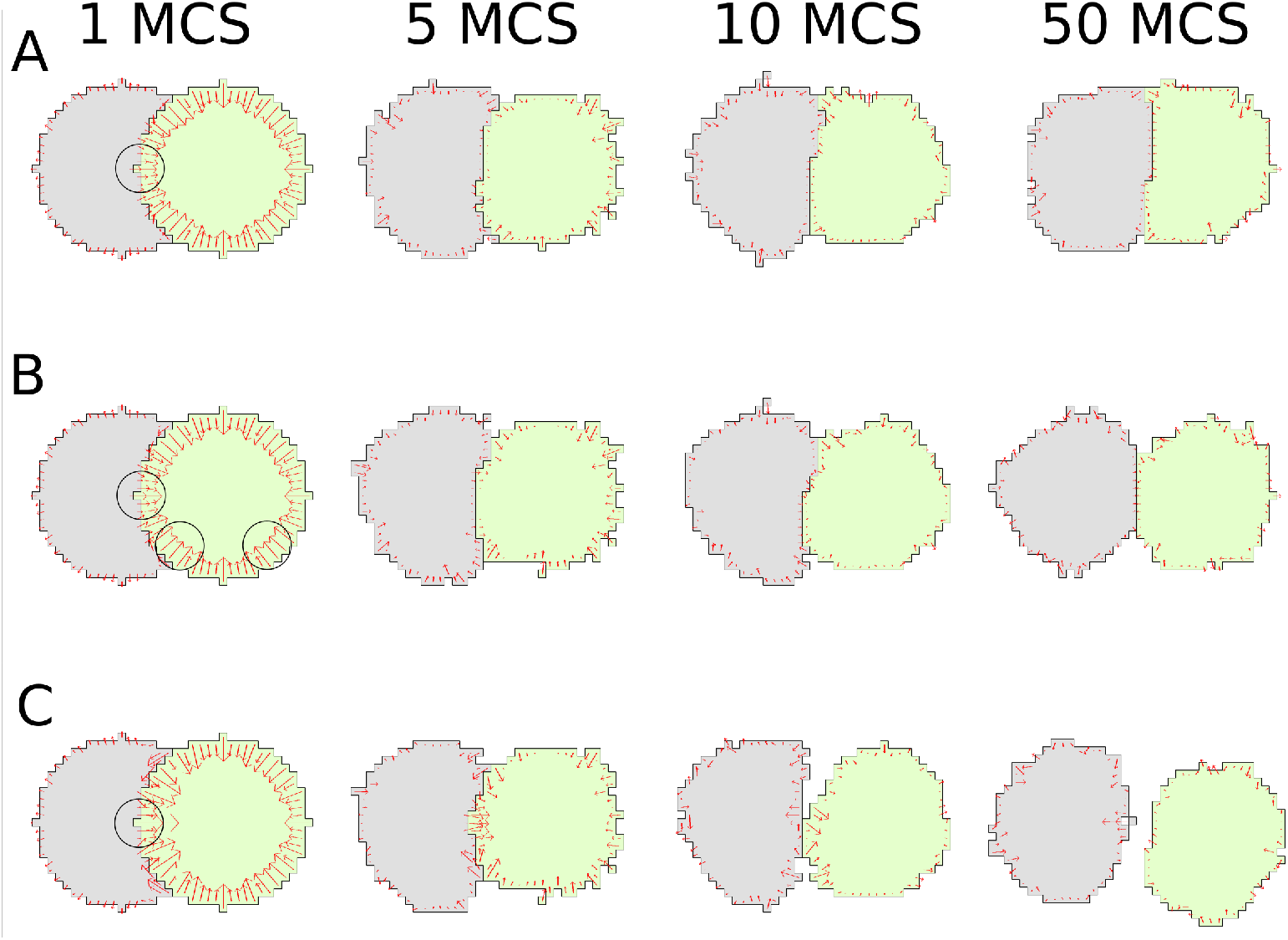
Forces due to cell-cell adhesion. Two CPM interacting cells in a time sequence from left to right. (A) cells adhere strongly *J*(1, 2) < 2*J*(0,1), (B) neutral adhesion of cells to medium and to one another *J*(1, 2) = 2*J*(0,1), (C) cells de-adhere, *J*(1, 2) > 2*J*(0,1); CPM parameters used were λ_*p*_ = 0, λ_*a*_ = 8, *a* = 300, *J*(1, 0) = 1800, *T* = 300. We used *J*(1, 2) = 1800 (for the adhesive), 3600 (for the neutral) or 7200 (for the repulsive) cases.

Comparing forces at cell-cell interfaces for the three adhesive energies (centered black circles at 1MCS), we find that the force at a cell-cell interface is lowest in A, and highest in C. This is consistent with Eq. (0.8).

### Two motile cells with internal signaling

We next asked how internal signaling in each of two interacting CPM cells would affect their mutual adhesive forces. To explore this question, we assumed the wave-pinning signaling in each of the cells, starting from initially cells with uniform signaling activities except for elevated activity along their left edges. Results are shown in Figure 8. The reaction-diffusion (WP) equations lead to rapid polarization of signal activity inside the cells, as before. High signal strength was associated with local contraction of the cell edge, and area constraint then led to net motion. The two moving cells maintained contact due to their assumed high adhesion (low energy of cell-cell interfaces). While initially, cells moved in roughly the same direction, at some later point, they started to rotate. This trend continued during the simulation. We show the internal signal distribution in 8A, the total force computed from the CPM Hamiltonian in 8B, and the active force due to the Rho-like signal in 8C. It is apparent from the latter that forces cause a torque, leading to the observed rotation.

**Figure 8.**
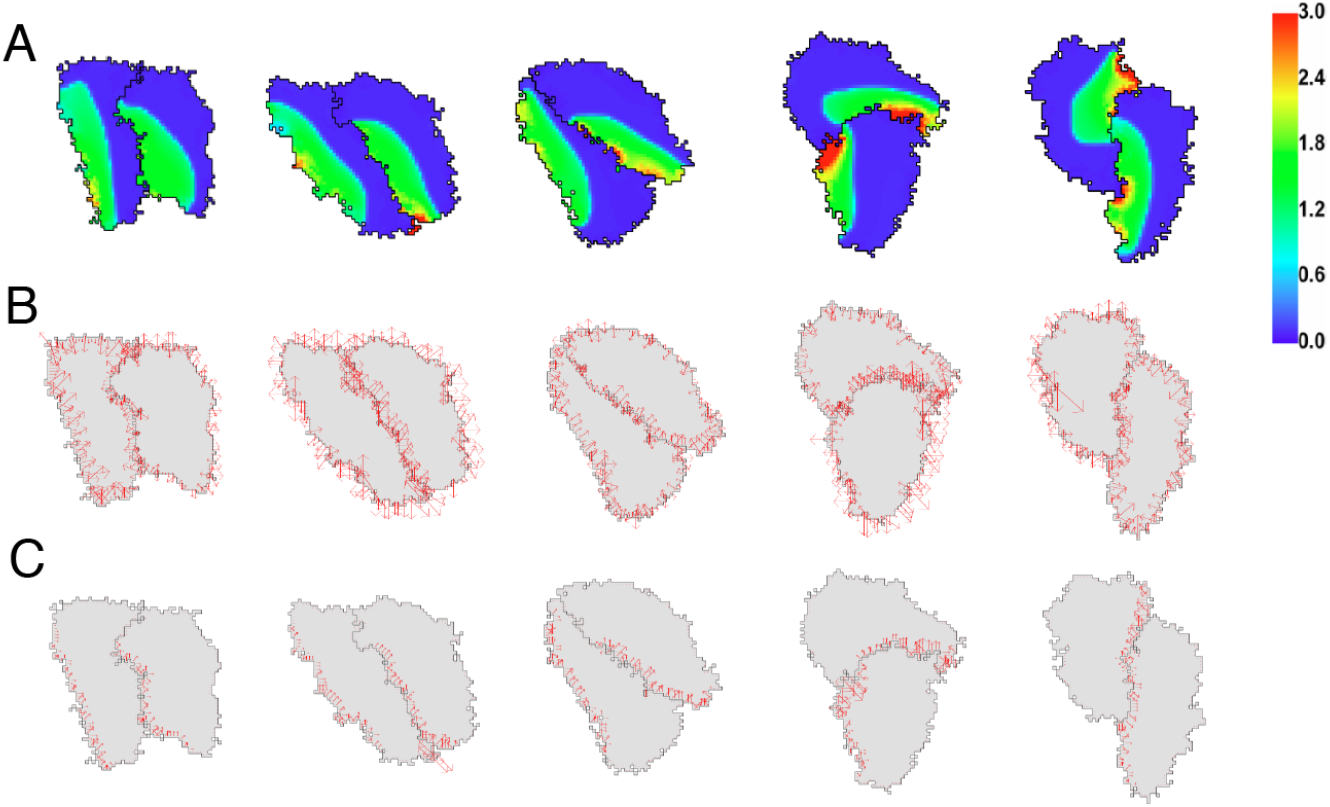
Edge forces in two adhering cells with internal signaling. (top) The level *u* of signaling activity, (middle row) total force exerted by each cell, (bottom row) mutual forces due to signaling contraction alone. The cells polarize and circulate about one another. Parameters for the CPM are λ_*a*_ = 10, *a* = 1000, λ_*p*_ = 0, *J*(1, 0) = 4000 = *J*(1, 2), *T* = 100, *β* = 20 (for dHp). Parameters for the reaction-diffusion system are provided in the Supplementary Information. High signaling activity (red) leads to local edge contraction. The configurations are shown at MCS 224, 375, 450, 525 and 600.

### Dynamic force fields in large multicellular aggregates

Finally, we sought to test our methods on simulations of larger cell aggregates. We asked whether the known dynamics of cell sorting, e.g. [19–22], would correlate well with force fields that can now be directly visualized. For this purpose, we adopted the cell-sorting benchmark test cases, where dynamics are well-established. That is, we considered three typical cases, with two cell types and three distinct relative heterotypic and homotypic adhesions, leading to the classic checkerboard, separation, and engulfment scenarios.

Figure 9 shows a time sequence of the model cell aggregate for the “separation” case. Initially, cells are randomly mixed. Here, *J*(*AA*) = *J* (*BB*) = 900, *J* (*AB*) = 9000 (where A are green and B are grey cells), so that a relatively high energetic cost results from interfaces of unlike cell types (heterotypic interfaces). This means that the adhesive forces between green and grey cells are high and repulsive. Evident from Figure 9 are high forces that build up at heterotypic interfaces. (See zoomed regions.) By Monte-Carlo step 400, we find regions where cells have separated. Cell boundaries continue to adjust for some time, accounting for fluctuations between outwards and inwards-pointing forces in a given cell during these transients. By 1000 MCS, many of the separated boundaries have equilibrated to a large extent, and localized forces on those boundaries have relaxed. A few remaining cells are still compressed or stretched away from their preferred rest area and perimeter, and are seen to experience significant forces. Later, (5000 MCS, Supplementary Figure 24) separated clusters round up. Interestingly, these static images, in combination with the force-map allow for easy visualization of parts of the aggregate that are still actively deforming.

**Figure 9.**
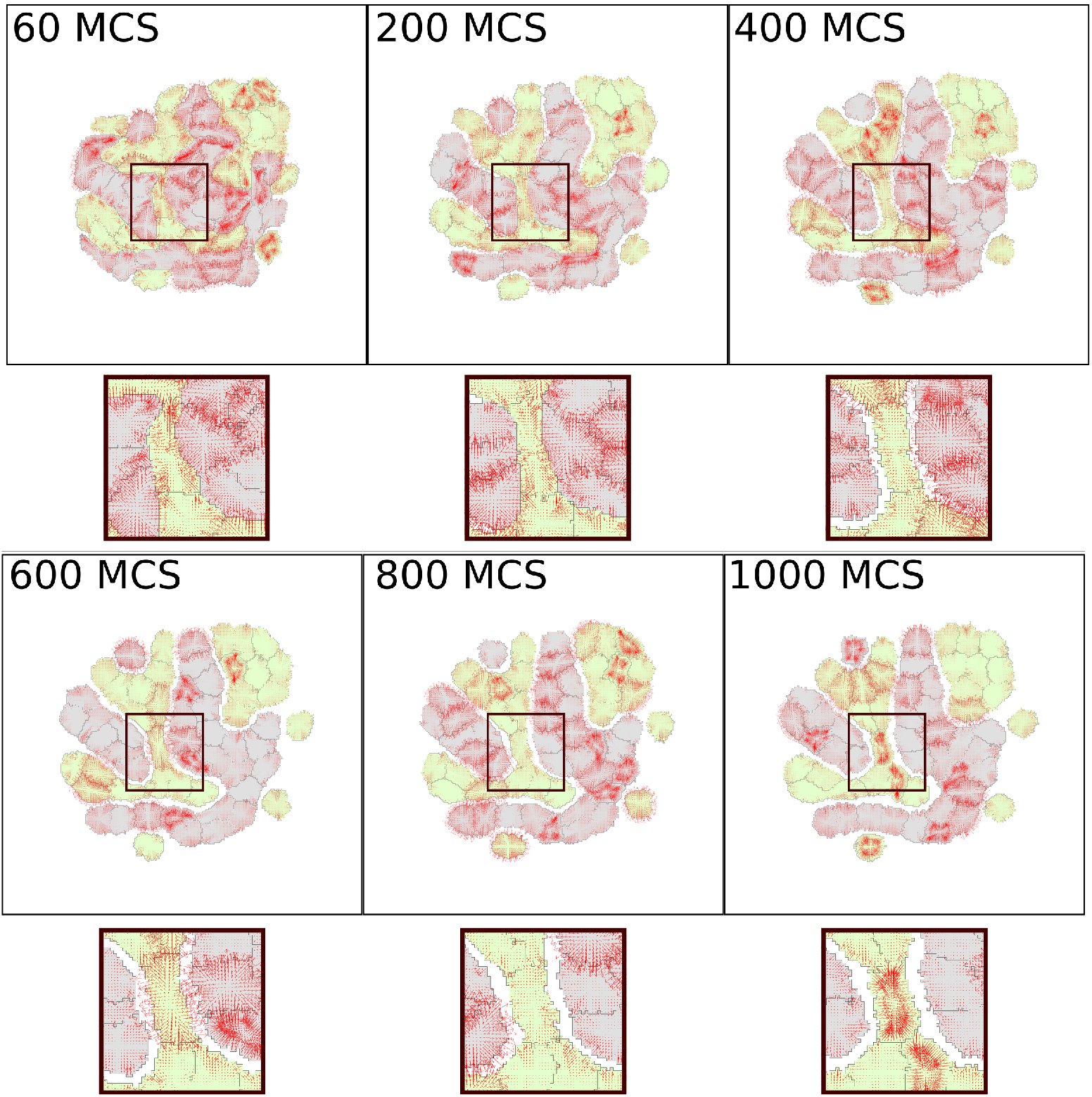
Separation cell sorting simulation with force visualization. Parameter values were *a* = 300, λ_*a*_ = 1000, *p* = 67, λ_*p*_ = 20, *J*(0, grey) = 1800, *J* (0, green) = 1800, *J* (grey, grey) = 900, *J* (green, green) = 900, *J* (grey, green) = 9000, *ξ*(*r*) = 18, and *r* = 3 for all neighborhood calculations. The cellular temperature *T* was set to 600.

We show two related scenarios in the Supplementary Information. A checkerboard cell-sorting case is illustrated in SI Figures 25 and 26. The engulfment case is shown in Figures 27 and 28.

## Discussion

Computational cell models promise to be a useful tool in testing hypotheses in single and collective cell behavior. A range of platforms is used, including force-based simulations of cells as self-propelled particles, points, spheres, or ellipsoids obeying Newtonian physics. A number of geometric cell models, including vertex-based (polygonal) cell simulations [23,24] are based on energy minimization [25,26] or on explicit springs and damping forces. One advantage of the Cellular Potts formalism, is that cell shape can be modeled in detail, is highly dynamic, and captures fluctuations seen in real cells.

Like any model, the CPM has its limitations, as described by [27]. Among these is the absence of an actual time scale in the “Monte Carlo Step”, mandating a definition of time scales by other methods (see, e.g., [2,28]). Here we have addressed a second common criticism of the CPM, namely that it bears no relationship to cell forces and mechanics. We have shown a direct link between Hamiltonian and corresponding forces.

Previous authors have combined classic CPM with external methods of tracking forces. Lemmon and Romer [16] assumed that a cell acts as a contractile unit resulting in a ‘first moment of area’ representation for the force distribution. Rens and Merks [3] adopted this same method. Such a model produces reasonably realistic force fields, but are not necessarily consistent with the CPM Hamiltonian, as these forces are assigned independently of the assumed form of *H*. Albert and Schwarz [5] adapted the CPM Hamiltonian to a form that had analytical expression for force on the cell edge. This was based on the curvature of the cell. They used a marching square algorithm to determine the curvature of the pixelated cell to calculate the cell edge force vectors and applied a smoother to distribute those forces in a region around the cell edge.

Magno et al. [2] used the link between forces and gradients of potential energy 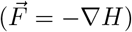 to write down the tension (*γ*), the pressure (Π), and the total force 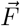 for the basic CPM Hamiltonian,

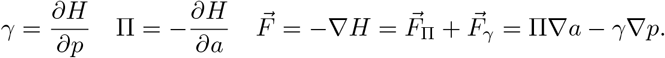

The authors used these relationships to derive a dynamical system for the size of a spherical cell, and to map cell size dynamics onto a 2-parameter plane with composite parameters. Our paper has taken motivation from their ideas to devise an algorithm for numerically computing forces directly from the CPM Hamiltonian, for an arbitrary cell shape, and for multiple cells.

Our numerical framework derives forces directly from small changes in CPM Hamiltonian when the cell configuration changes by a small “spin flip”. While the CPM Hamiltonian predicts forces at cell edges, simple interpolation and smoothing to decrease grid effects were adopted. We showed that this approximation for the forces gives reasonable results for a range of cell shapes. Importantly, the most basic CPM Hamiltonian reproduces forces that are qualitatively consistent with experimental data. Our algorithm applies not only to single cells but also to multicellular simulations. The computed force-fields provide insights to cell deformations accompanying three typical cell sorting experiments, where most but not all cells equilibrate with their neighbors. In such simulations, force fields within the clusters help track and understand the global and local dynamics of the cell collective. From the force-fields we can appreciate simulated cell motions and a more tangible connection between the Hamiltonian and cell behavior.

The approach is an approximation and has limitations that we summarize here. First, the classic Hamiltonian approximates a cell as an elastic element tending to retract/expand towards a specified rest area and rest-length circumference, which is a grossly simplified view of a cell. Moreover, in matching CPM predictions to data, we find multiple sets of CPM parameters that give rise to very similar qualitative agreement. Improved calibration of Hamiltonian to cells of given type would require more specific data and are beyond the scope of this paper.

A second issue is that the CPM Hamiltonian only changes for cell edge displacements, and so, only prescribes a force-field that is restricted to the cell edge pixels. We have assumed simple interpolation, with zero force at the cell centroid, but this is, to some extent, arbitrary. As seen in the data in Fig. 6, a polarized cell can have multiple points at which the force vanishes. Hence, the internal force field should not be over-interpreted. A second limitation is the attribution of forces to cell shape alone, neglecting active and heterogeneous structures (stress fibers, focal adhesions, cytoskeleton anisotropy, etc.) that make each cell a “living machine”. There are generalizations to our idea that could improve on this simplification. In previous work, Maree et al [7] assembled a more detailed internal signaling CPM model for a single motile cell that included actin filament orientation and pushing barbed ends (regulated by active Cdc42, and Rac), as well as edge contraction (due to GTPase Rho, as in our simple examples in Figs. 4 and 8). Such details can be added for greater consistency with motile cells. Alternately, inclusion of other Hamiltonian terms such as directional polarity (Mareé and Grieneisen, unpublished), or heterogenous, space-dependent, Hamiltonian terms could be used to generalize these ideas. Finally, in principle, internal cell structures (nucleus, focal adhesions) could be added, each having distinct properties (i.e. values of parameters λ_*a*_, λ_*p*_, *J* etc.) This would necessitate a refinement of the CPM accordingly.

An issue with CPM computations is that Monte Carlo steps are not scaled to actual time. This issue is generally resolved by scaling the motion or cell cycle of CPM cells to real cell speeds or cycle times. Based on typical cell size, typical forces cells produce, and typical values of viscosity, one could also use the relationship *v* ≈ *F/ξ* to devise a time scale. See also [28] for a proposed definition of the time-scale.

We have carried out partial validation of the method against single-cell data. Traction force microscopy has also been used to quantify patterns of stress in multicellular aggregates [18]. Stress is usually localized at the periphery of a cluster of cells, while at cell-cell interfaces the stress is lower [29], suggesting that the cluster acts as a single contractile unit. Inside the cluster, forces are highly dynamic and localized forces can occur due to cell proliferation or rotating motion [30]. Great progress has been made in visualizing force fields, and it is likely that modeling and computation will contribute to an understanding of how traction force are precisely regulated and what are consequences of the force dynamics on single and collective cell behavior.

## Materials and Methods

### Cellular Potts Model

In a Cellular Potts model (CPM), each “cell” consists of *N* lattice site, assigned a unique index (“spin number”). Parts of the domain containing no cells are indexed 0 by convention. A spin flip copies the spin value of a source lattice site 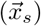 to a target site 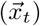, typically in a Moore neighborhood (one of eight nearest-neighbor pixels). The configuration change 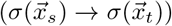 results in a change Δ*H*, in the Hamiltonian.

Our Hamiltonian is given by Eq. (0.2) Many “spin flips” are attempted, but a each is accepted with probability

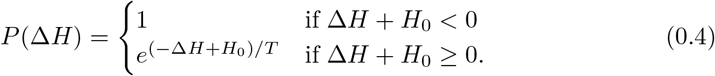

where the “temperature” *T* ≥ 0 governs the magnitude of random fluctuations and *H*_0_, is a yield energy to be overcome. (Typically *H*_0_ = 0.) The CPM favors changes that decrease the energy of the configuration, while allowing fluctuations.

### Approximating forces at points along cell boundaries

We discretize the gradient of the Hamiltonian, from Eq. (0.3) as follows. Let *h* = Δ_*x*_ = Δ_*y*_ be the given grid size in 2D. For each point 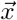 on the border of a cell of configuration *σ*, consider a small local change, protrusion or retraction (Fig. 1). The local “spin flip” at 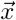 produce a small change in the Hamiltonian. We can compute the force components 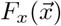 and 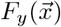 at 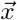 using a centered difference approximation to the first partial derivative (accurate to 2nd order):

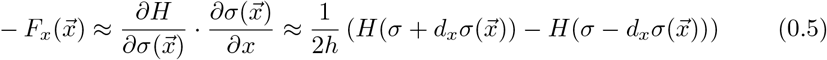

and similarly for the component 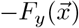.

This recipe defines forces at each point along the boundary of each isolated CPM cell. In the Supplementary Information we discuss other simplification and special cases. We implement steps to (1) improve accuracy, and reduce grid effects (2) interpolate boundary forces to the cell interior (in 2D) (3) Generalize the idea to multiple cells and (4) Compare to measured force fields for real cells.

### Reducing the grid effects in perimeter calculations

Because cell boundaries are pixelated, a grid-effect is introduced in the above calculations (Supplementary Figure 14). We reduce this artifact using enhanced CPM neighborhood calculations inspired by [2]. Briefly, at each boundary site we define a weighted average of forces with weights given by “local cell perimeter” as computed using neighborhood summation. (We use a neighborhood radius *r* = 3.) We find that this correction results in forces that are roughly orthogonal to the (refined) cell boundary. In the Supplementary Information, we provide details and discuss how accuracy is affected by neighborhood radius.

### Phenomenological force fields in the interior

The methods described so far only provide a representation of the force field associated with the cell perimeter. We use simple interpolation from boundary sites to a point in the cell interior, typically the centroid of the region. This phenomenological choice, following [3–5], leads to a 2D force field.

### Intracellular reaction-diffusion system and protrusive forces

To model intracellular signaling that affects cell shape, we implement the wave-pinning reaction-diffusion (RD) model of [6] in the 2D cell interior, and compute the evolution of the RD system with no-flux boundary conditions at the evolving cell boundary. Methods for our numerical computation, analogous to those of [7] are described in the Supplementary Information. To link the internal chemical profile to forces on the cell boundary, we assume a Rho-like edge contractility: the “Rho activity”, *u*, close to the cell edge, is assumed to augment the local Hamiltonian changes by additional terms dH of the form ±*βu* for protrusions/retractions. In this way, the distribution of *u* can locally affect the probability of movement of the cell edge. After the cell edge moves, *u* is redistributed locally to avoid numerical mass loss, as described in the Supplementary Information.

### Comparison with experimental data

We obtained traction force microscopy (TFM) data from Jocelyn Etienne and Claude Verdier for two cancer cell lines (T24 and RT112) as described by [8]. The authors plated cells on polyacrylamide gels containing fluorescent beads, and computed traction forces from bead displacements and known gel rheology [9]. We interpolated from their triangular to our rectangular grid (Supplementary Figure 17), and optimized the CPM parameters with respect to data at one time point using a Latin Hypercube sampling method (see Supplementary Information and Tables 1 and 2). The CPM and data force fields are then displayed on the same grid, and their difference, directional deviation, and relative magnitudes are computed and displayed for comparative purposes.

### Generalization to multiple cells

For a system of multiple cells, we decompose the total Hamiltonian into contributions *H^i^* made by each cell,

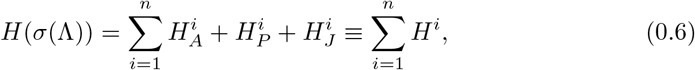

where 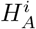 and 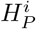 are as in Eq. (0.2) for cell *i* and where *H_J_* is generalized to accommodate cell-cell adhesion energies,

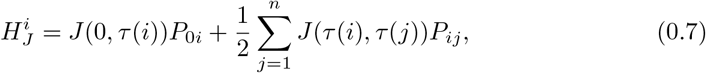

where *n* is the number of cells, *τ*(*σ*) the cell type of cell *σ*, *P*_0*i*_ is the boundary length of cell *i* in contact with the medium and *P_ij_* is the length of the cell *i*- cell *j* interface. (The factor 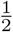 corrects for double-counting of each interface.) The finite difference computation of forces along interfaces then follows from the single cell case. (See also the Supplementary Information).

It has been shown in other papers (see, e.g. [5]) that force exerted by each cell can be reduced to the form

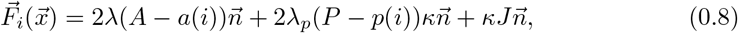

where *κ* is the curvature, 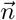 is the unit normal vector, and *J* is either *J*(0,1) or *J*(*i, j*)/2.

## Supplementary figure captions

**Supplementary Fig. 10.**
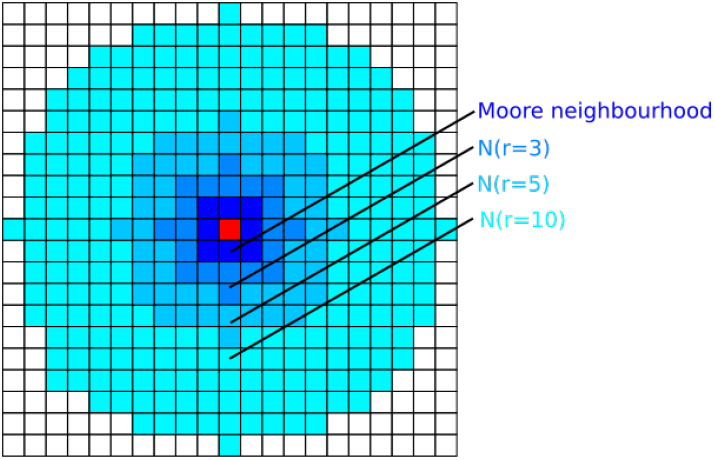
Neighborhoods 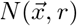 of various orders with radii *r* = 3, 5,10 around a lattice site 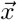 (shown in red).

**Supplementary Fig. 11.**
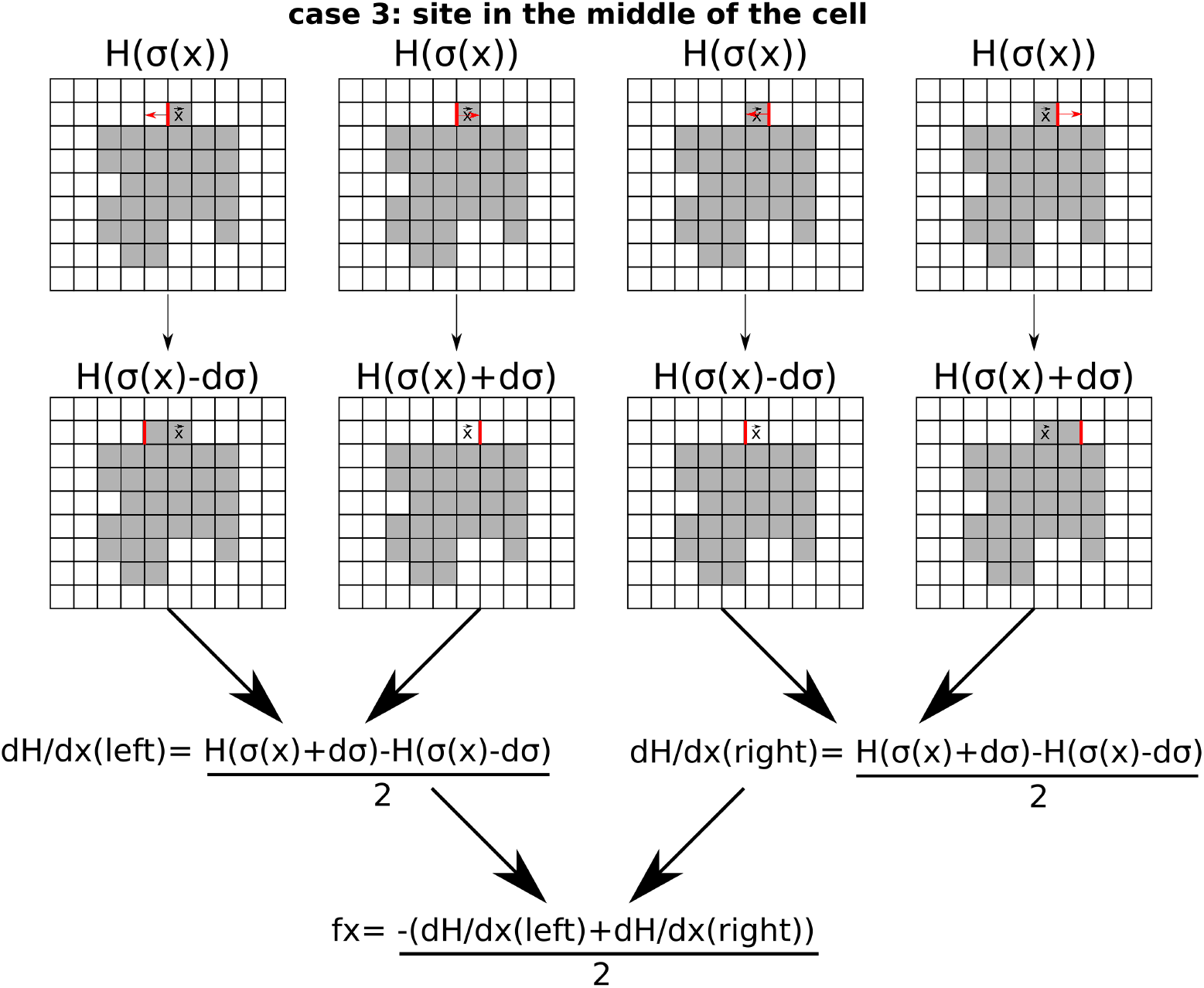
Examples of four possible spin flips used to compute *F_x_* based on Eq. (0.5).

**Supplementary Fig. 12.**
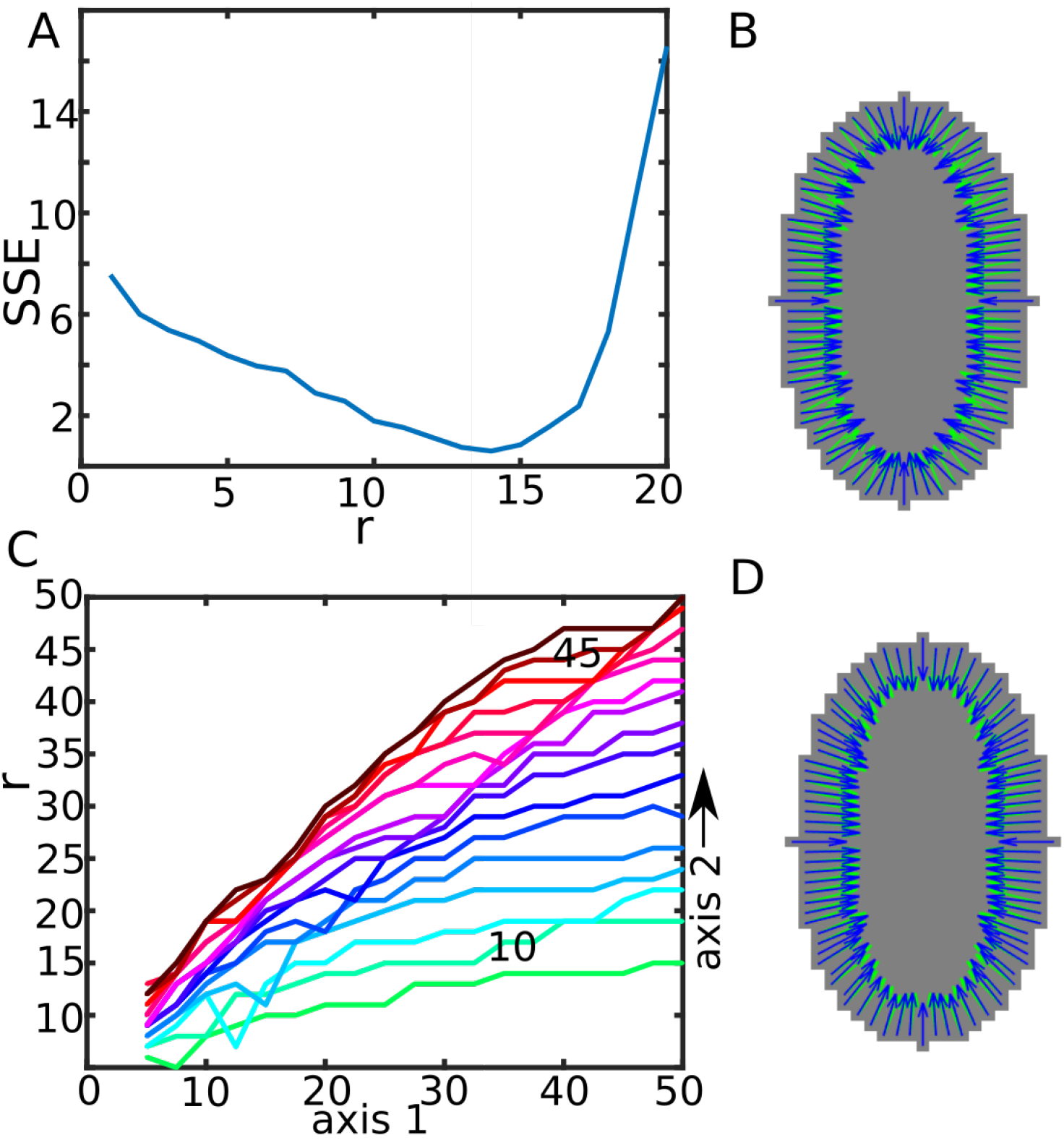
The effect of the neighborhood radius used for smoothing the cell boundary forces for ellipsoidal cells. (A) Sum of square errors (SSE) between normalized CPM force vectors and true unit normal vector (–*b* cos(*θ*), –*a* sin(*θ*)) to an ellipse with axes 10 and 20. The error is minimized at *r* = 14. (B) True normal vectors (green) to an ellipse with axis 10 and 20, compared to CPM forces smoothed with radius *r* = 3 (blue). (C) Neighborhood radii *r* corresponding to minimal SSE for ellipses with various axes lengths between 5 and 50. (D) Same as (B) but with smoothing radius *r* = 14.

**Supplementary Fig. 13.**
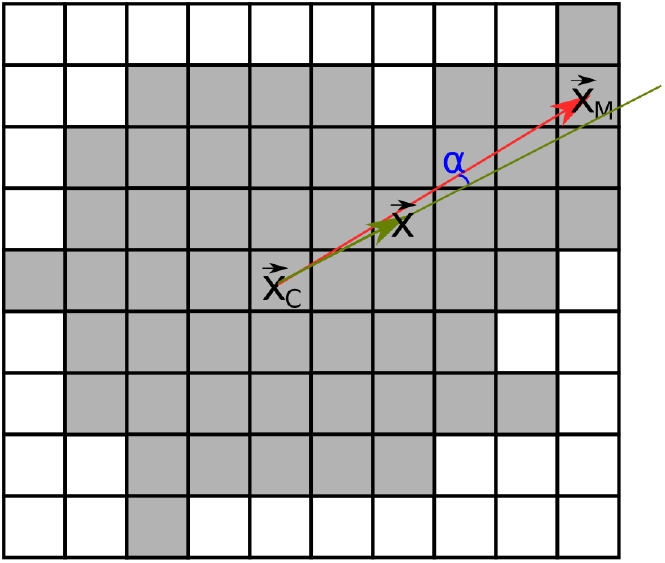
Interpolation used to compute force at a site 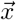 inside a CPM cell based on the centroid 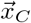 and the force predicted by the CPM at a boundary site 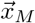 along the ray connecting the centroid and the given site. The ray was determined by minimizing *α* in Eq. 0.22.

**Supplementary Fig. 14.**
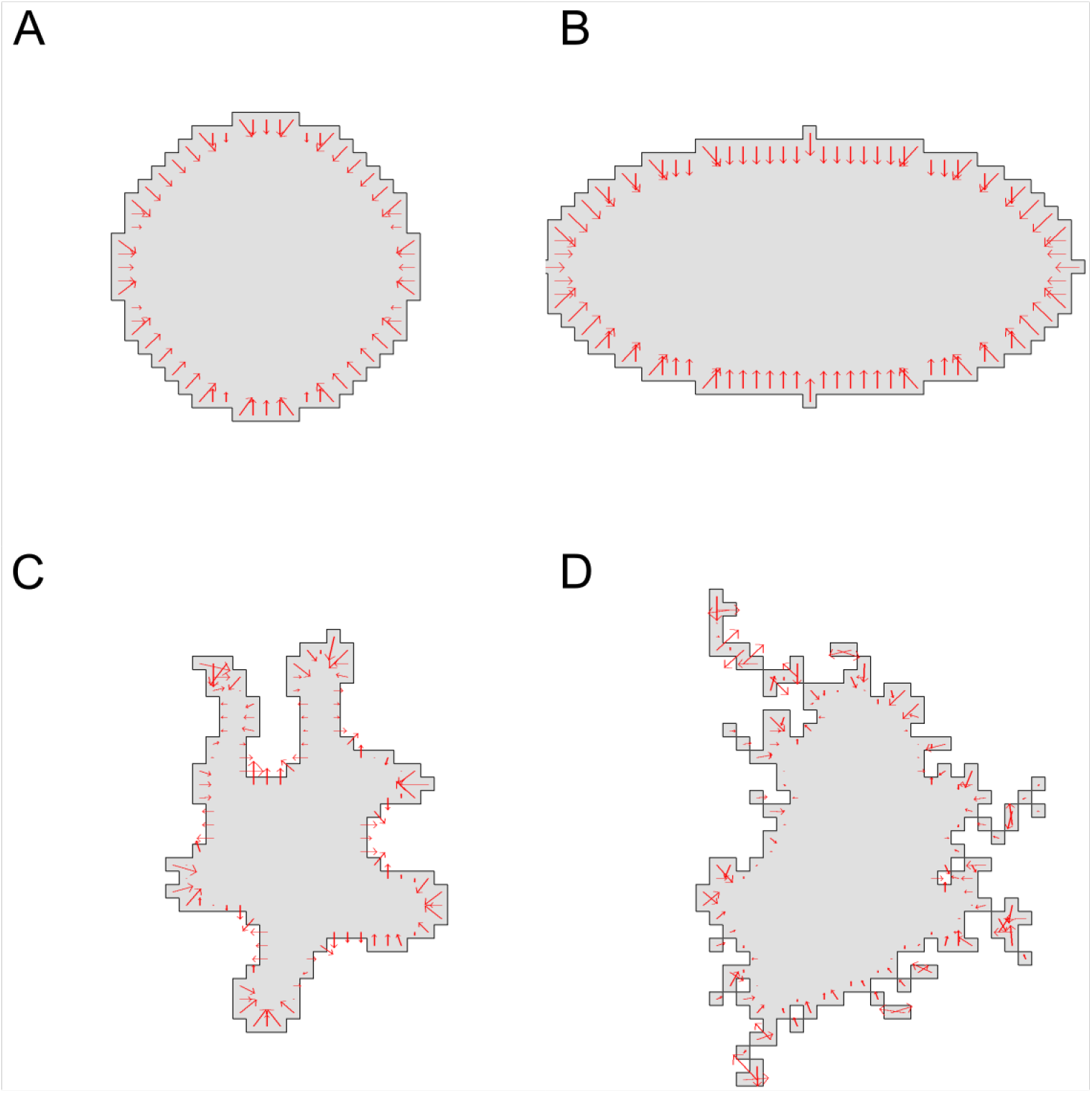
Cell edge forces for various simulated cell shapes in the CPM using only the finite difference approach, with no smoothing. (A) A circular cell with an area of 401, perimeter of 74, and a diameter of 23. (B) An elliptical cell with an area of 629, perimeter 101, and short and long axis 21 and 41. (C) An irregular shape with area 301 and perimeter 118. (D) A highly irregular cell shape with area 400 and perimeter 146. Parameter values were *a* = 300, λ_*a*_ = 10, *p* = 100, λ_*p*_ = 10, *J*(0,1) = 3000, *ξ*(*r*) = 18, and *r* = 3 for all neighborhood calculations. We used a grid of 50 by 50 lattice sites with Δ*x*=1.

**Supplementary Fig. 15.**
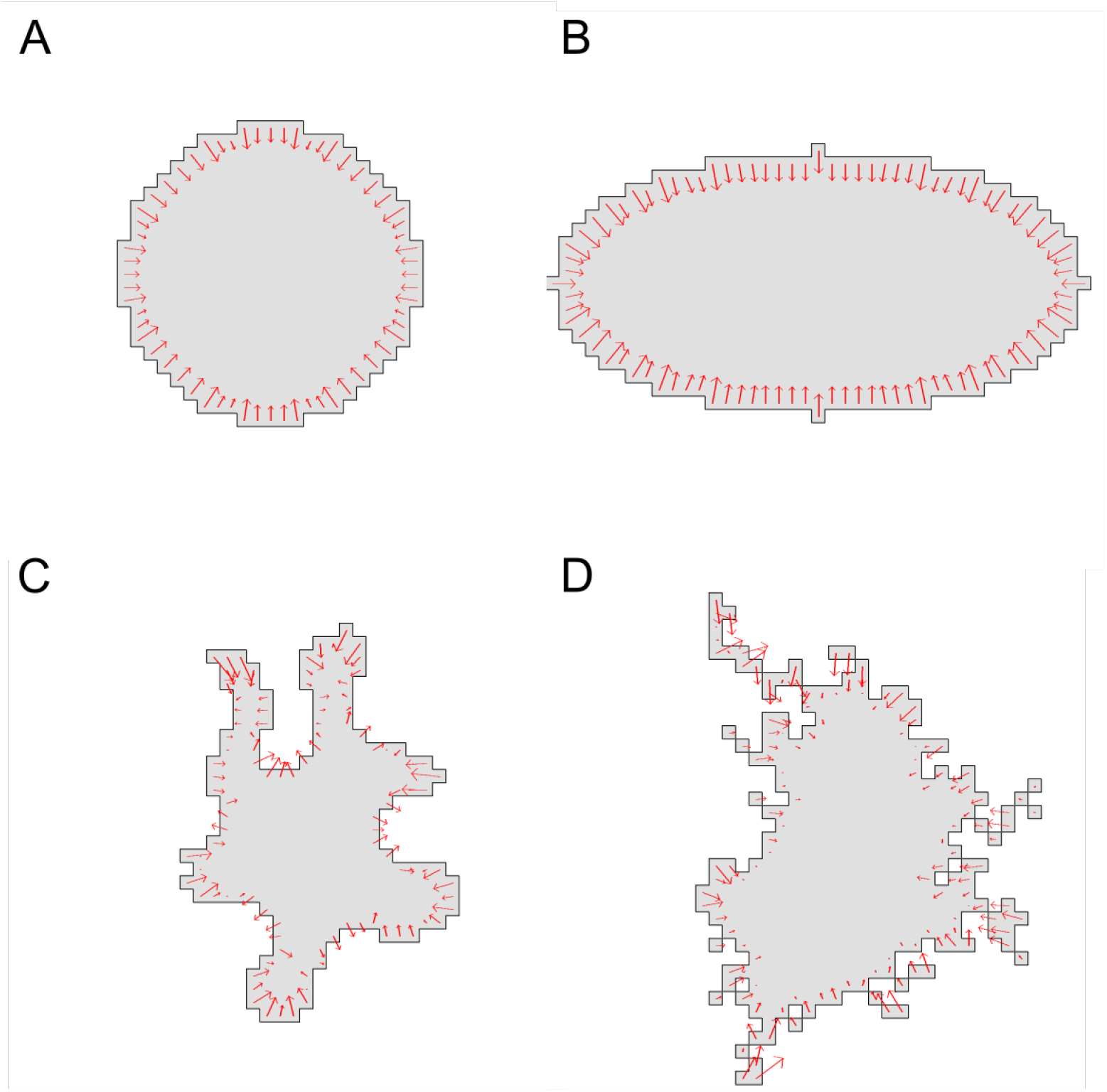
As in Supplementary Fig. 14, but with smoothing applied to the boundary forces. We used *r* = 3 for all neighborhood calculations.

**Supplementary Fig. 16.**
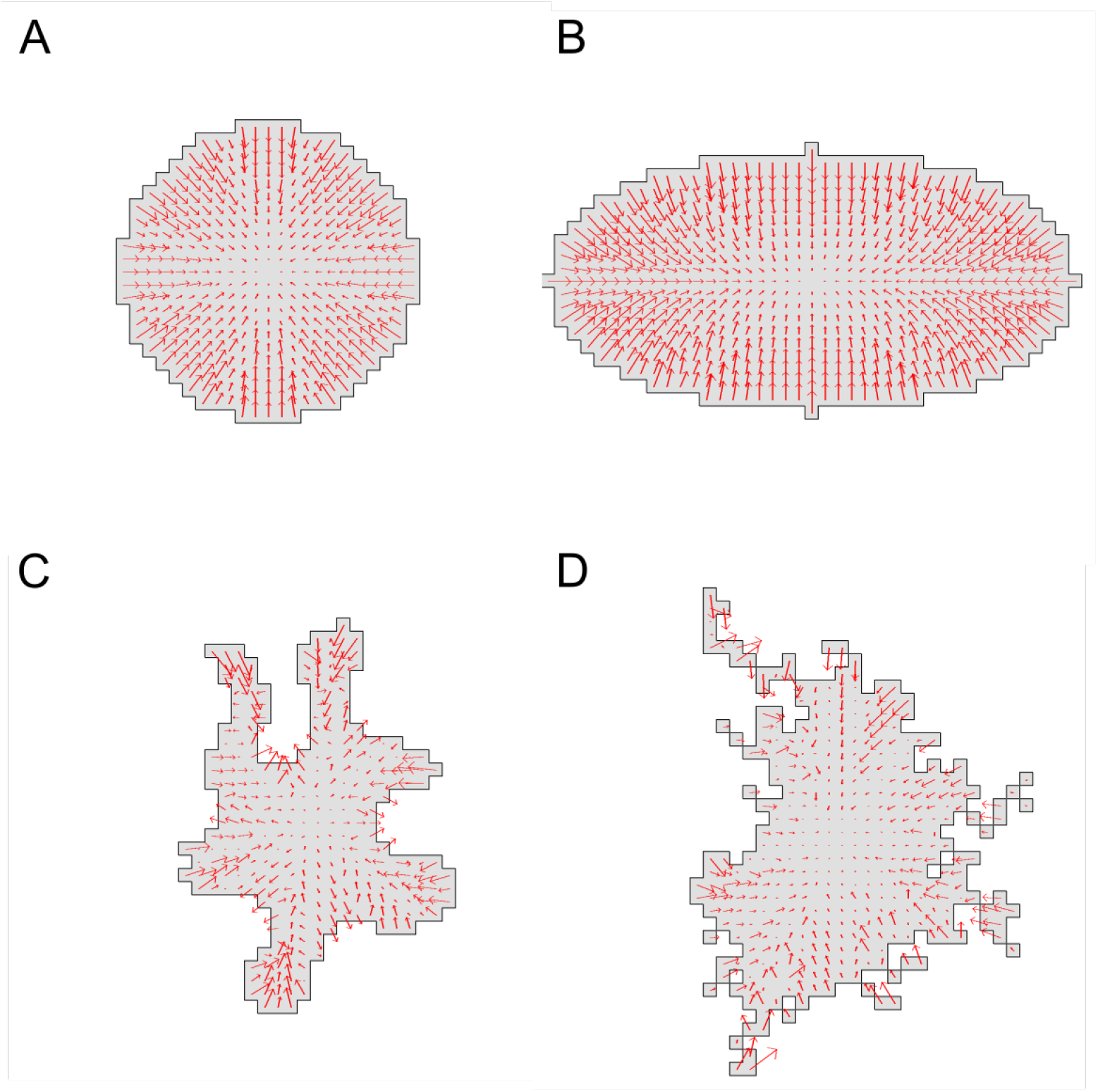
Interior forces computed with no smoothing for the cell shapes shown in Supplementary Fig 14.

**Supplementary Fig. 17.**
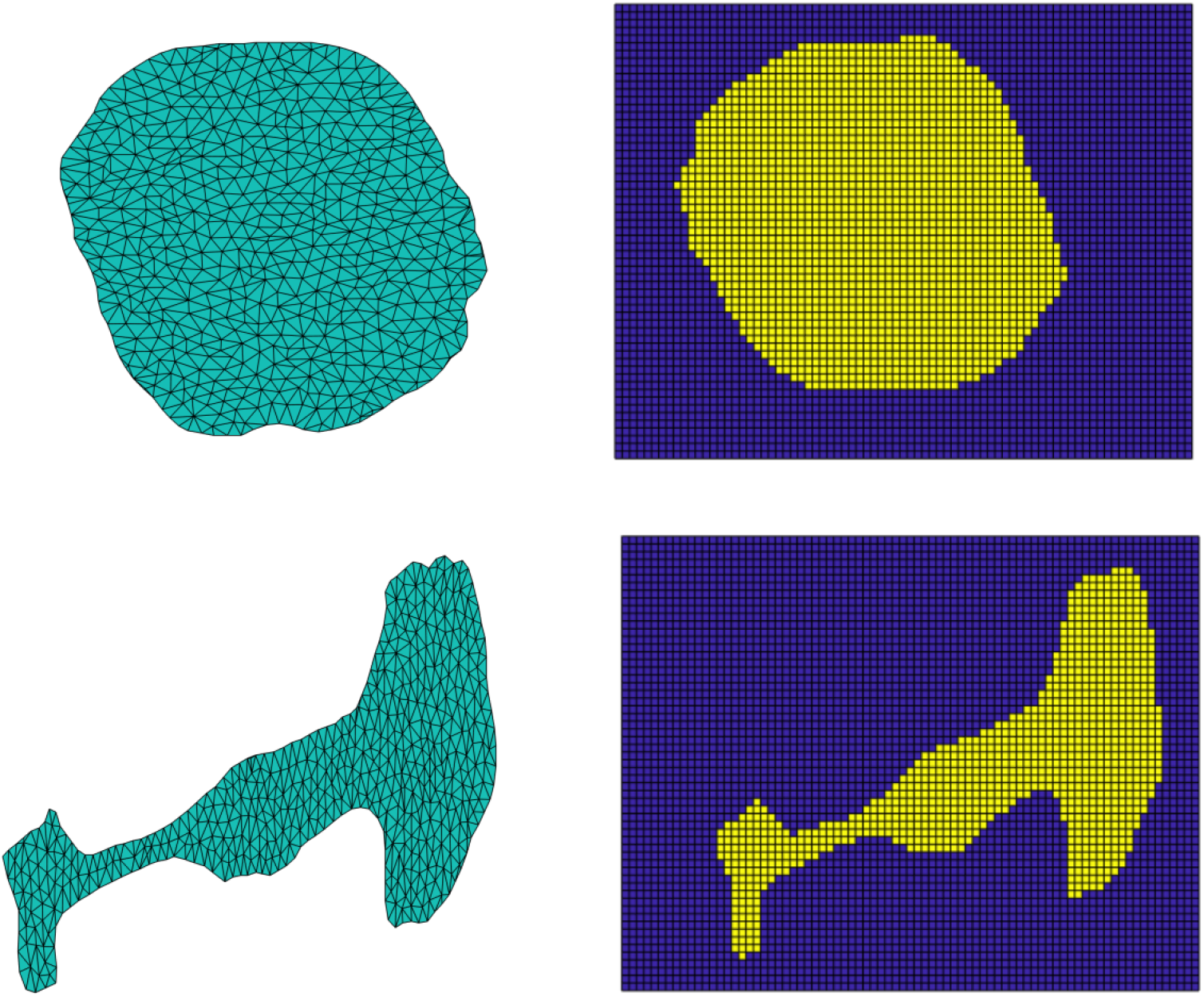
Triangular mesh on which cell traction data from [8] was supplied, and the corresponding CPM cell (spin value = 1).

**Supplementary Fig. 18.**
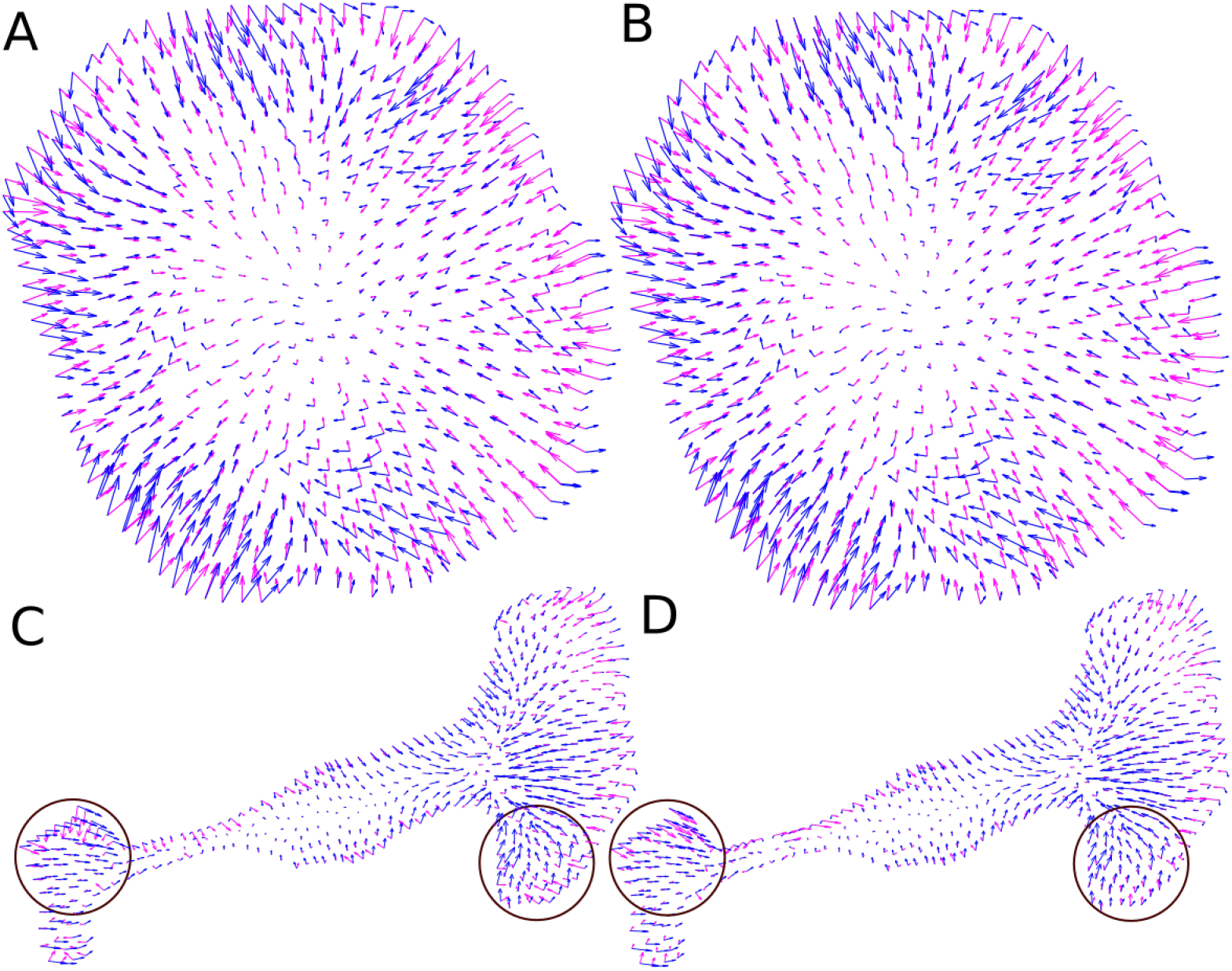
Force fields from data (blue) and CPM (magenta) using initial arbitrary CPM parameters for the round cell (A-B) and polarized cell (C-D). Radius of smoothing used was (A,B) *r* = 3, (C, D) *r* = 10. Regions of large deviation are circled.

**Supplementary Fig. 19.**
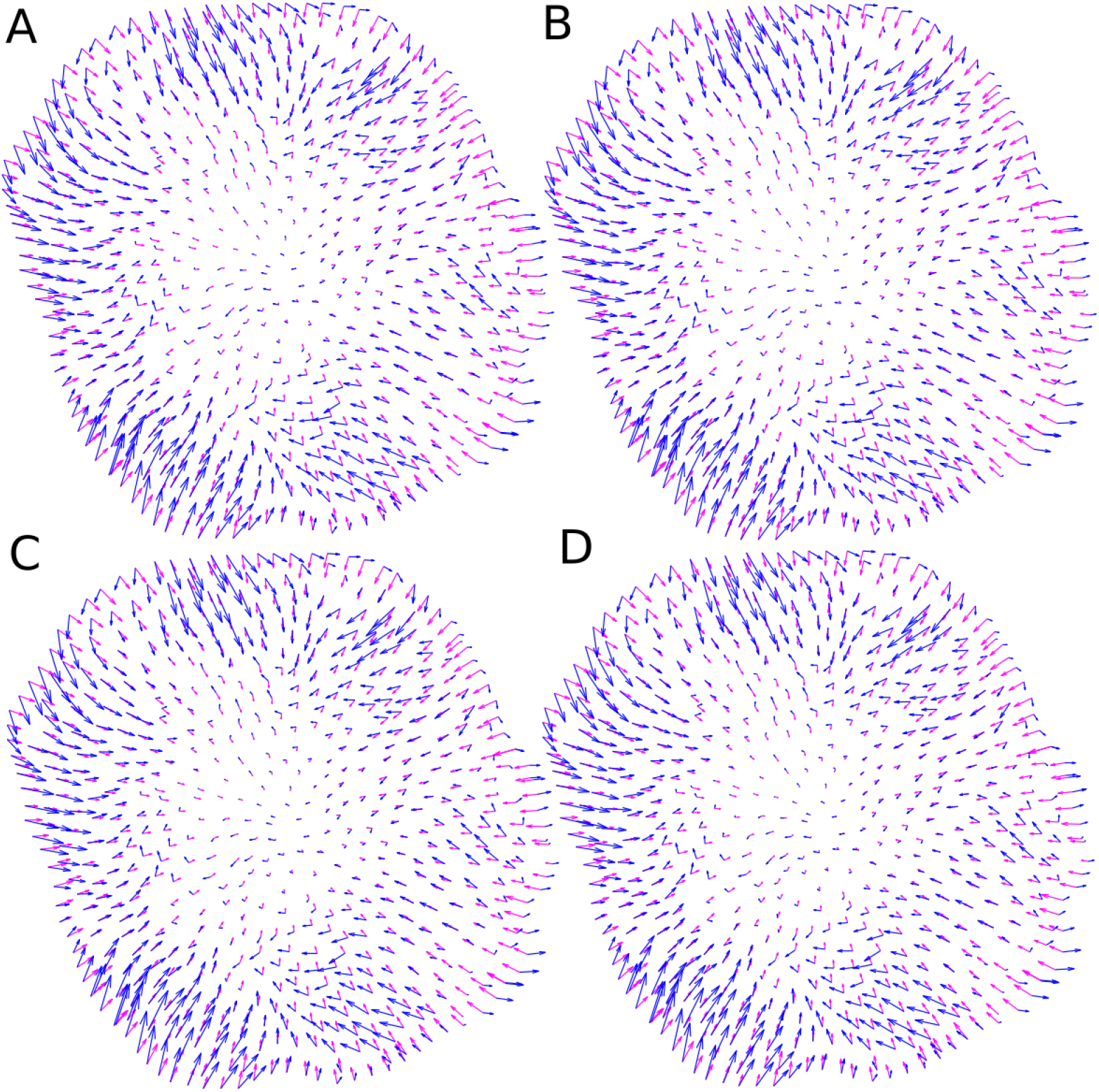
Fitting CPM parameters: Data (blue) and CPM (magenta) force fields for the round cell using the second (A), third (B), fourth (C) and fifth (D) best CPM parameter values. Parameter values are given in Table 1.

**Supplementary Fig. 20.**
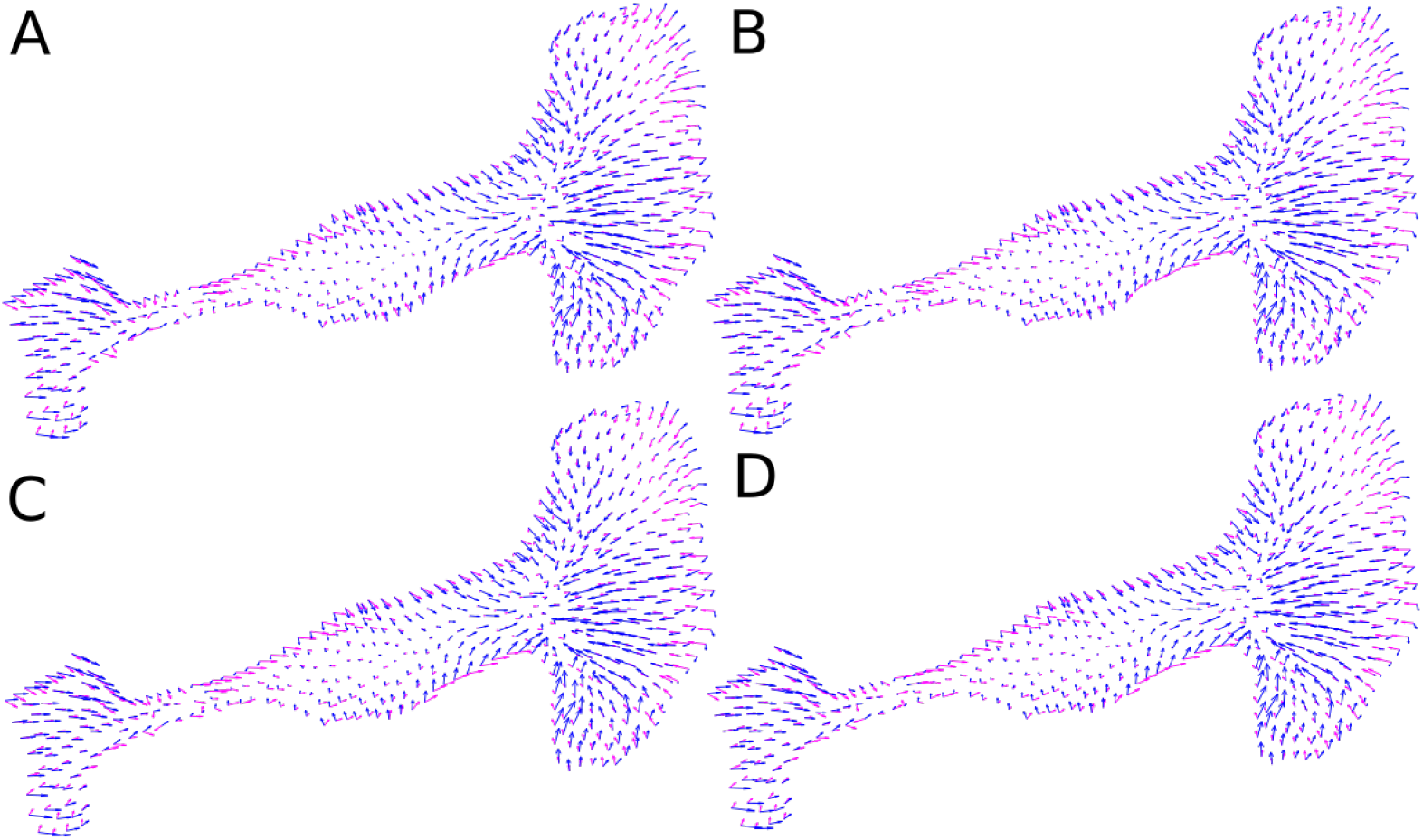
As in Fig. 19 but for the polarized cell using the second (A), third (B), fourth (C) and fifth (D) best CPM parameter values in Table 2

**Supplementary Fig. 21.**
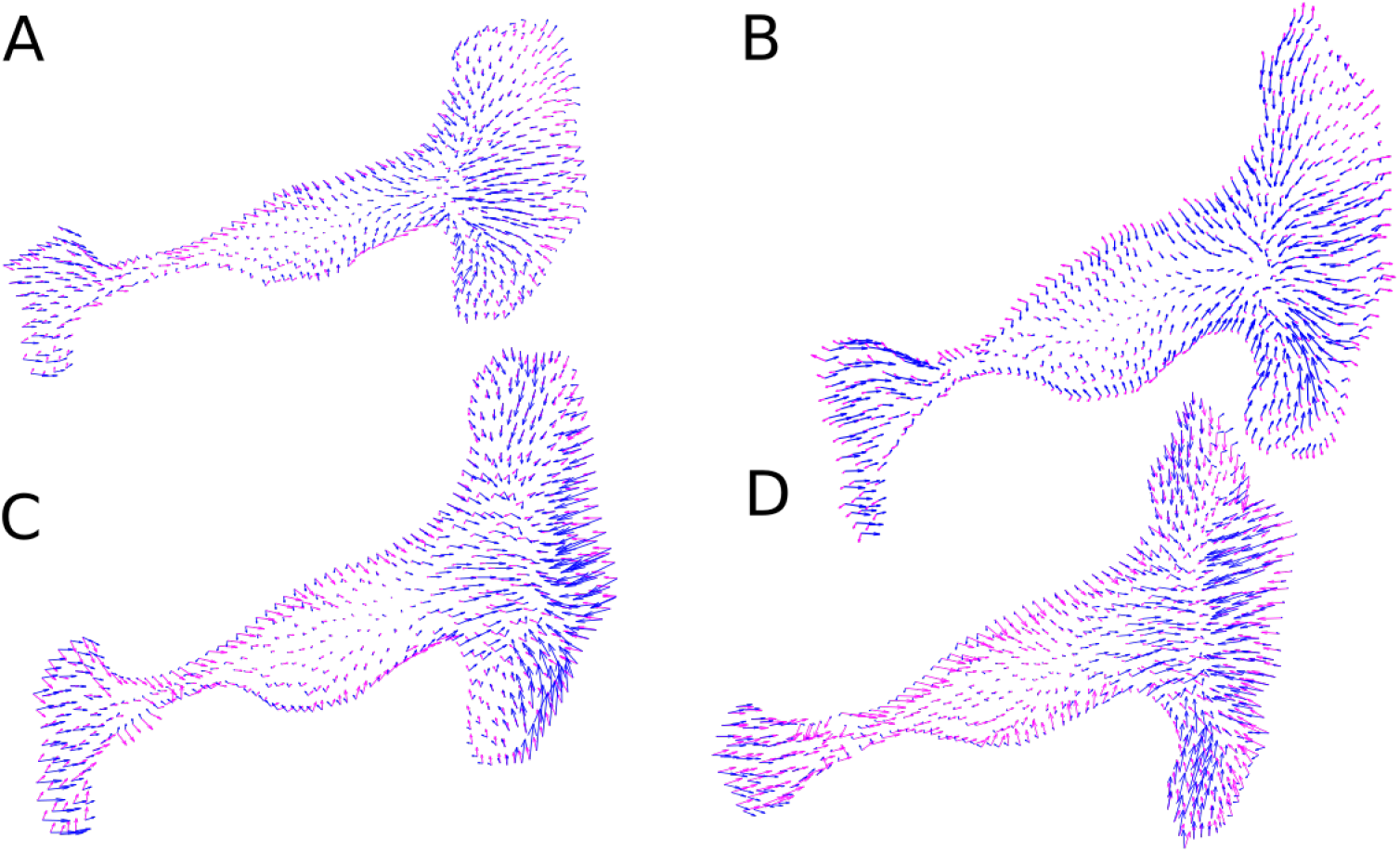
A time-lapse of cell motion and force fields from [8] showing data (blue) and CPM (magenta) force fields. The CPM parameters were as in Fig. 6 and row 1 of Table 2.

**Supplementary Fig. 22.**
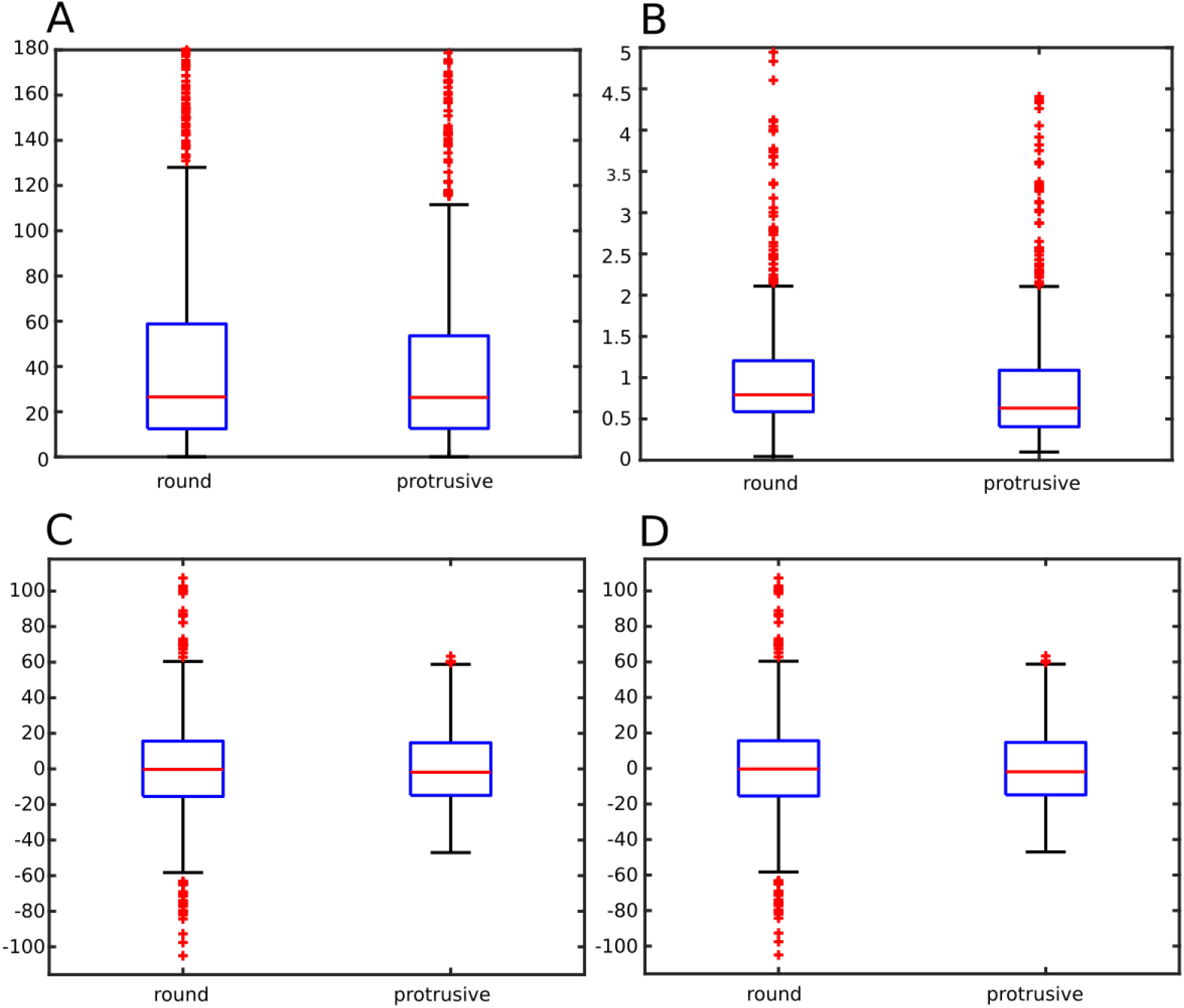
Correspondence between data and CPM predicted forces. Boxplots showing distributions of (A) the directional deviation (angle between experimental and model forces), (B) relative magnitudes of forces (C) deviation of x components and (D) y components of the forces.

**Supplementary Fig. 23.**
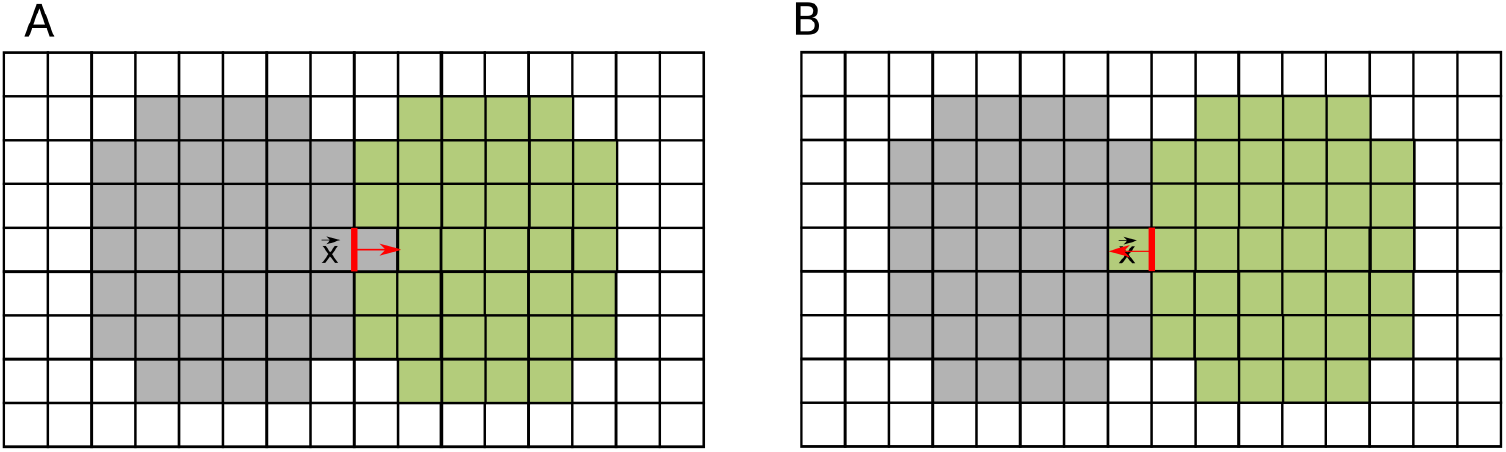
Multiple cells: Spin-flips used to approximate the force exerted by the grey cell at cell-cell interfaces (A) CPM spin-flip modeling extension of the grey cell, shifting the cell-cell interface to the right (B) CPM spin-flip modeling a retraction of the grey cell, shifting the cell-cell interface to the left.

**Supplementary Fig. 24.**
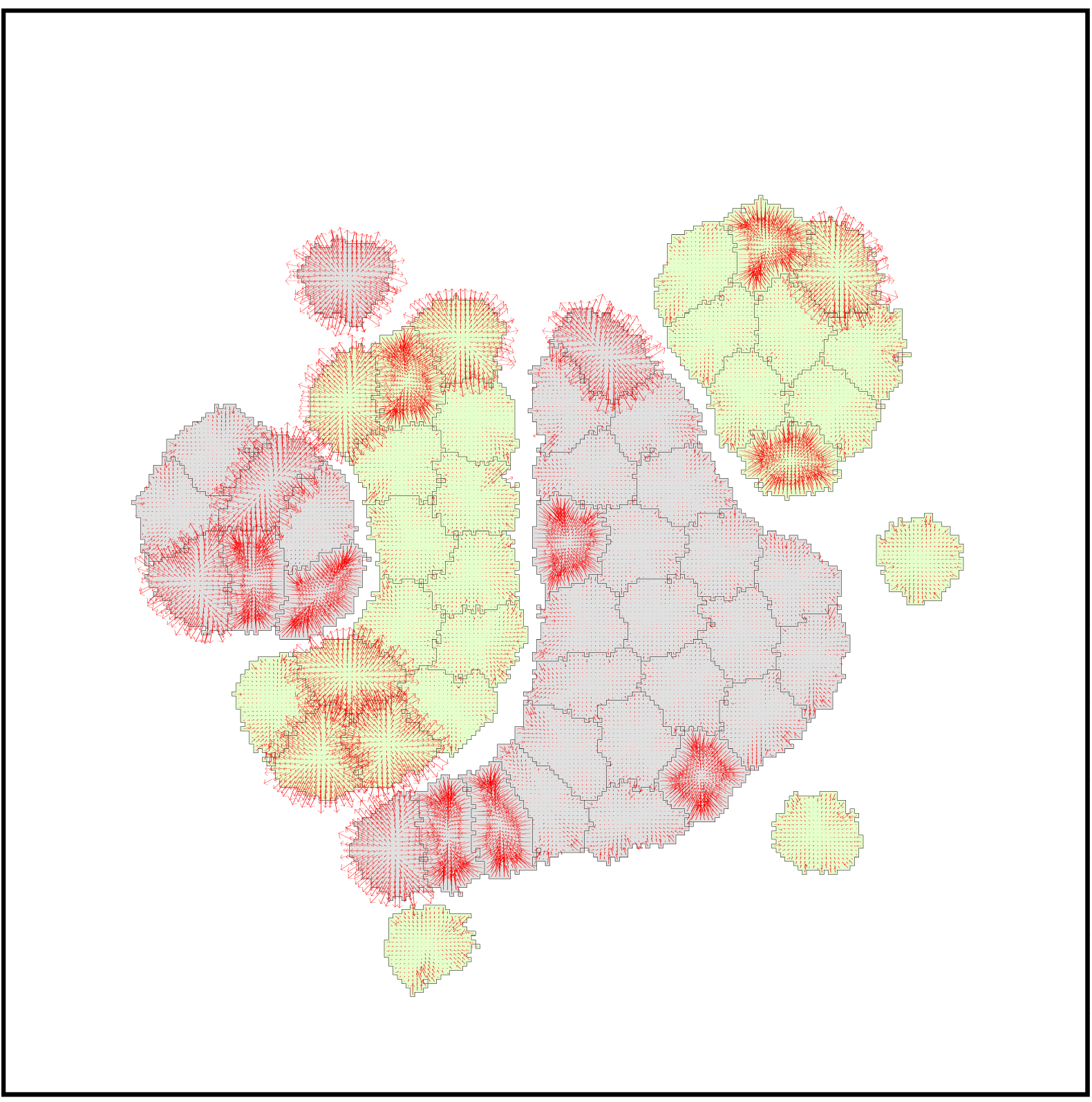
A separation cell-sorting simulation at 5000 MCS. Parameter values were *a* = 300, λ_*a*_ = 1000, *p* = 67, λ_*p*_ = 20, *J*(0, grey) = 1800, *J* (0, green) = 1800, *J* (grey, grey) = *J* (green, green) = 900, *J* (grey, green) = 9000, *ξ*(*r*) = 18, and *r* = 3 for all neighborhood calculations. The cellular temperature *T* was set to 600. Some cells are still experiencing large forces.

**Supplementary Fig. 25.**
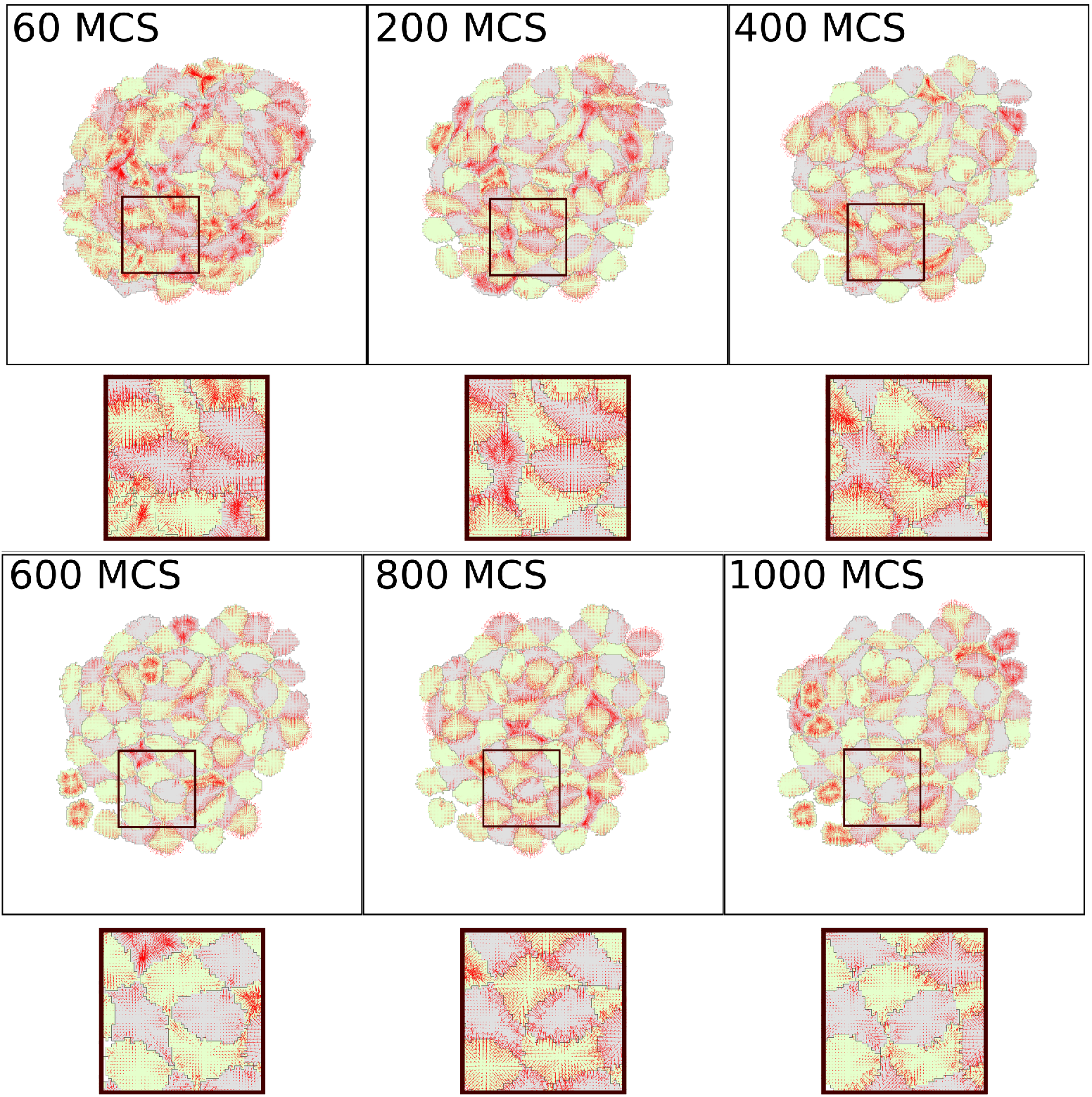
Checkerboard simulation with force visualization. Parameter values were as in Supplementary Fig. 24 but with *J*(grey, grey) = 7200, *J* (green, green) = 7200, *J* (grey, green) = 1800.

**Supplementary Fig. 26.**
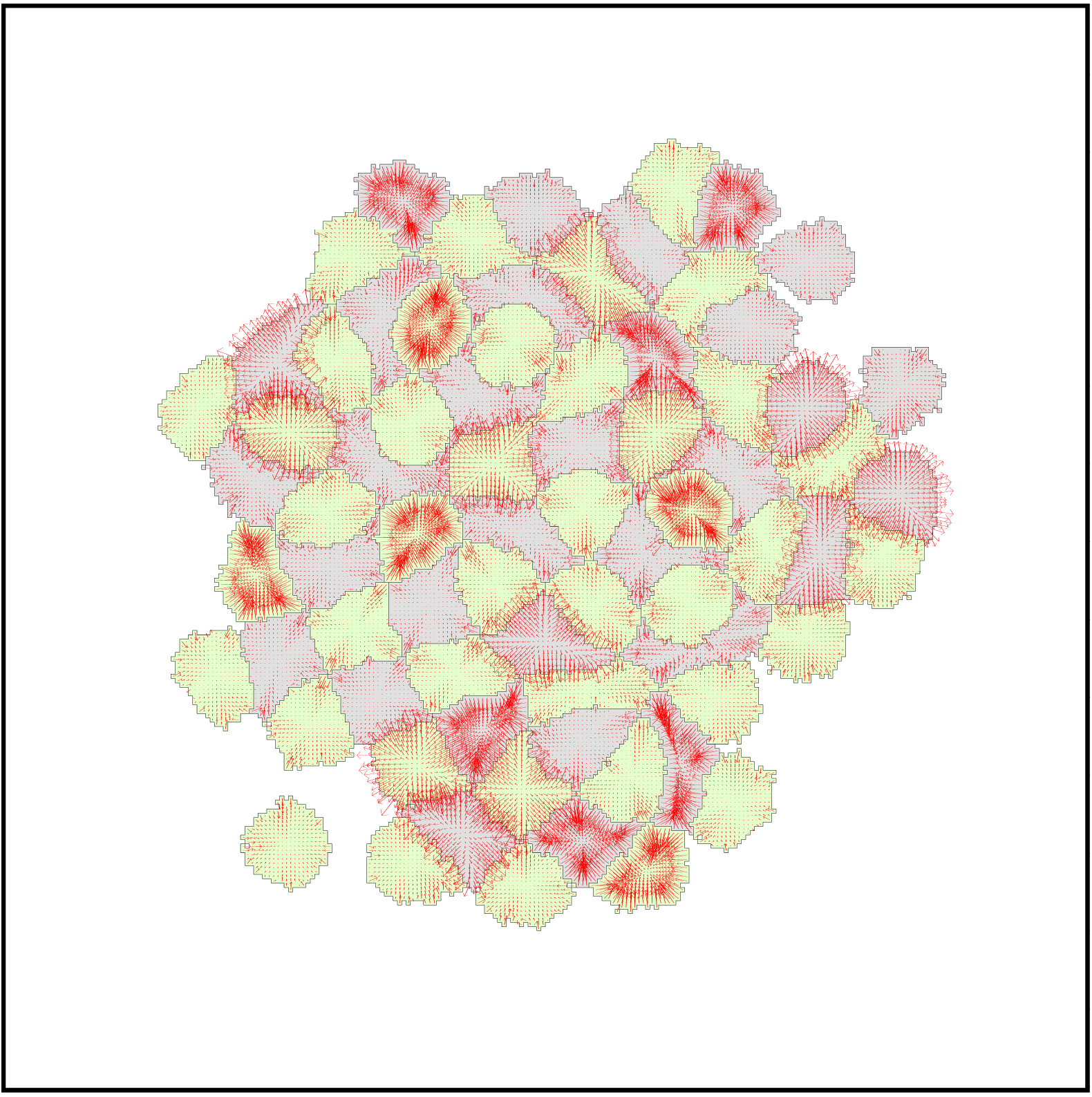
Checkerboard cell-sorting simulation at 5000 MCS. Parameter values were a = 300, λ_*a*_ = 1000, *p* = 67, λ_*p*_ = 20, *J*(0, grey) = 1800, *J* (0, green) = 1800, *J* (grey, grey) = 7200, *J* (green, green) = 7200, *J*(grey, green) = 1800, *ξ*(*r*) = 18, and *r* = 3 for all neighborhood calculations. The cellular temperature *T* was set to 600.

**Supplementary Fig. 27.**
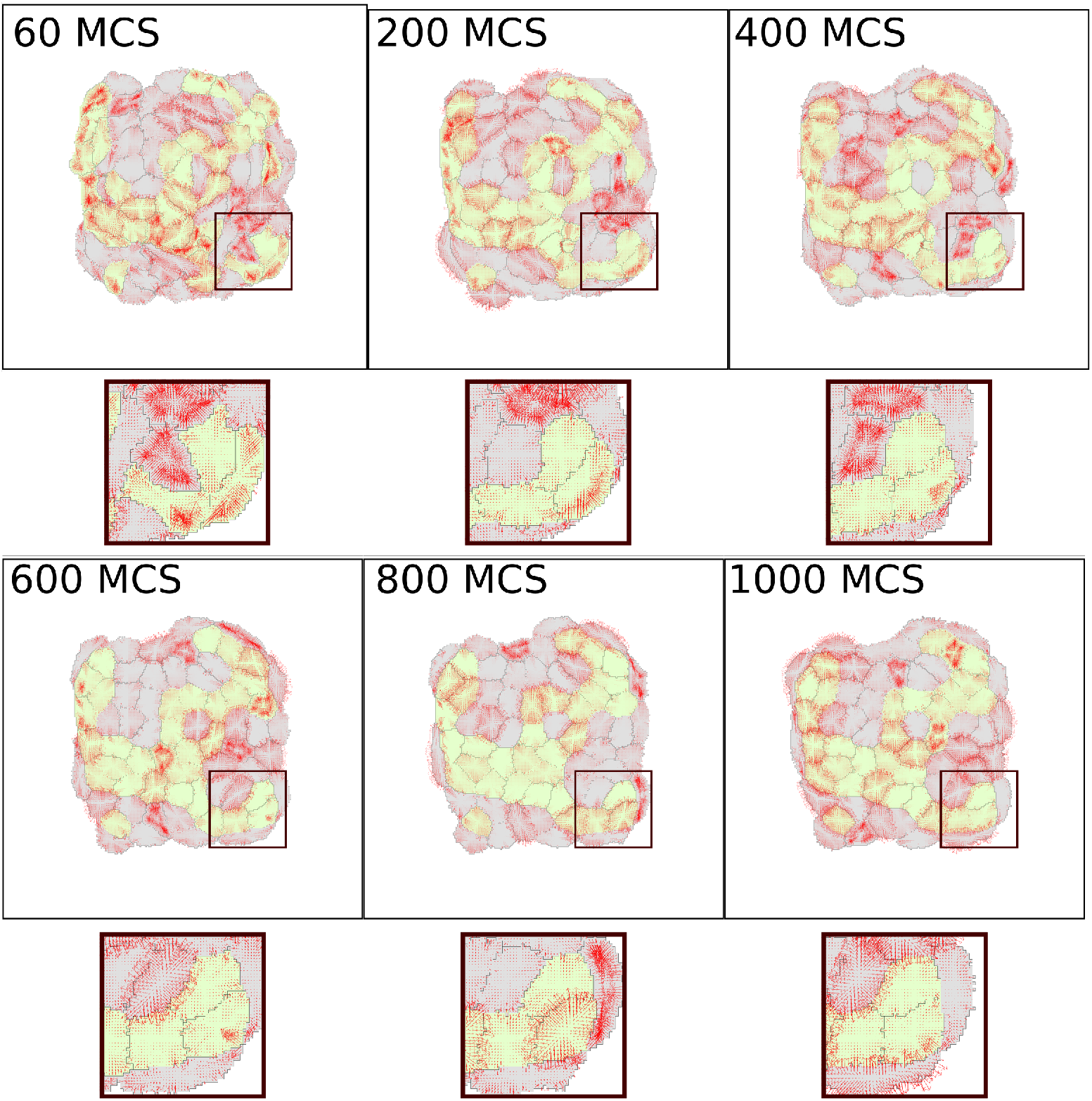
Engulfment cell-sorting simulation with force visualization. Parameter values were *a* = 300, λ_*a*_ = 1000, *p* = 67, λ_*p*_ = 20, *J* (0, grey) = 1800, *J* (0, green) = 9000, *J* (grey, grey) = 1800, *J*(green, green) = 1800, *J*(grey, green) = 3600, *ξ*(*r*) = 18, and *r* = 3 for all neighborhood calculations. The cellular temperature *T* was set to 600.

**Supplementary Fig. 28.**
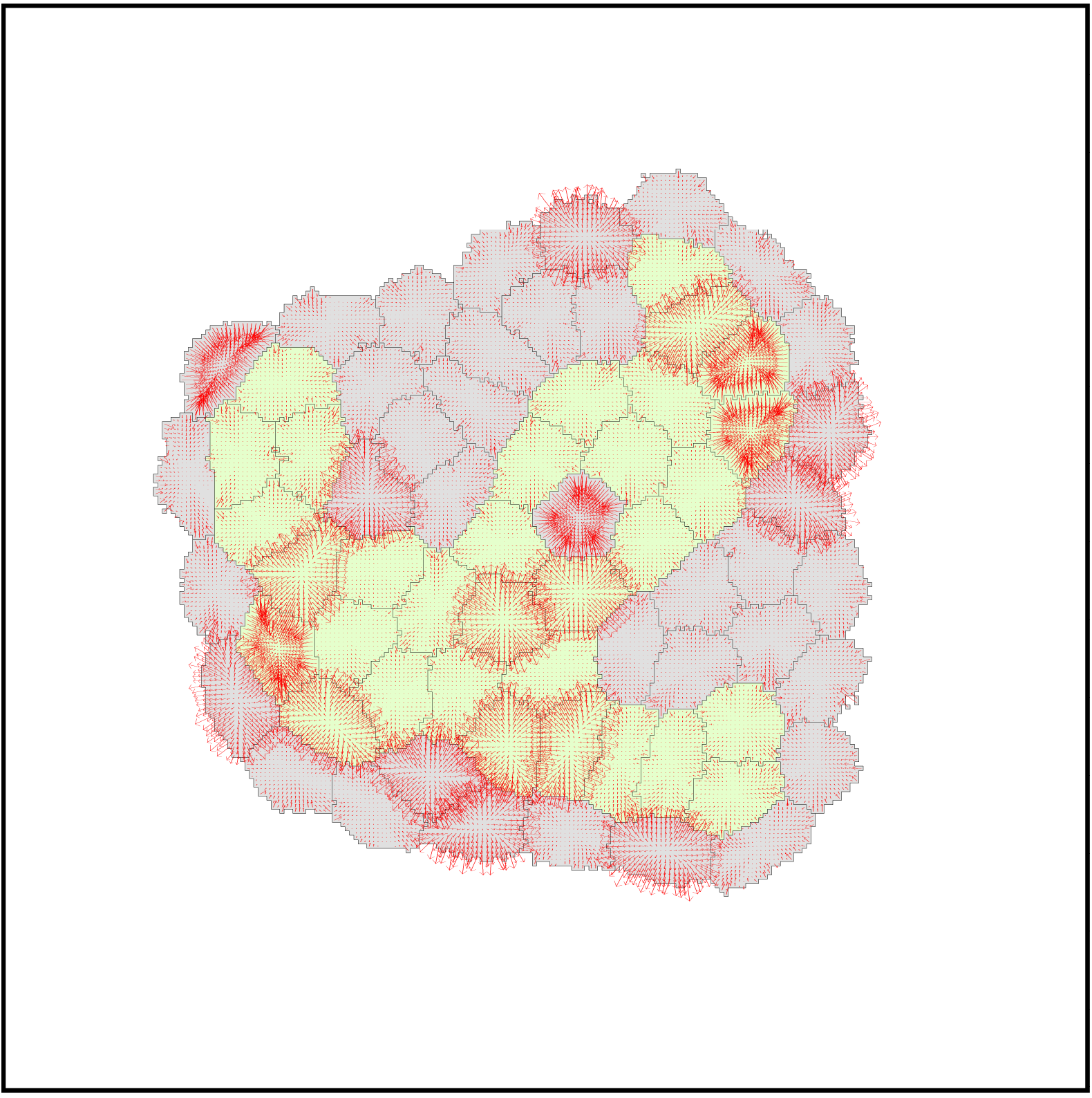
Engulfment cell-sorting simulation at 5000 MCS. Parameter values as in Supplementary Fig. 27.

## Acknowledgments

We thank Prof. Claude Verdier and Prof. Jocelyn Etienne for supplying original traction force microscopy data. We are grateful to Zachary Pellegrin (USRA student at UBC, summer 2018) for developing a reaction-diffusion solver on a deformable domain. We thank the Feng-Keshet group members for helpful discussions. This research is supported by an NSERC Discovery grant (41870) to LEK.

## Supplementary Information

### Cellular Potts Model (CPM)

In the CPM, cell shape is described on a discrete lattice and evolves through minimization of a Hamiltonian, analogous to an energy. Dynamic changes in cell shapes, interfaces, and positions result in changes in the Hamiltonian. The dynamics of the system are governed by the general principle of energy minimization, allowing for fluctuations that are akin to “thermal noise”. The latter helps to avoid getting trapped in local energy minima. The Hamiltonian is minimized by a Metropolis type algorithm, where cell edge movements are iteratively attempted and actually carried out if the movement decreases the Hamiltonian. For surveys of CPM and its applications, see [2,10].

In the CPM, both shapes and positions of each “cell” (configuration denoted by σ) evolve in time. Changes that minimize the Hamiltonian are favored. At each simulation step (Monte Carlo Step, MCS) every boundary pixel of each cell may “protrude” or “retract”. (Formally, these changes are denoted “spin-flips”, each corresponding to copying the spin value of a source lattice site (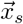: source site) onto a neighboring target lattice site (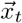: target site; 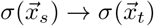). The target sites are typically the eight nearest-neighbor pixels of the source site (Moore neighborhood). While many such small spin-flips are tested, those that are actuated depend on the resulting changes in the Hamiltonian (Δ*H*) The probability of the move is assigned by Eq. (0.4), A common issue raised in the literature is that CPM simulations are not in correspondence with Newtonian forces, and hence non-physical. However, as argued eloquently in [2], the energy-based CPM formalism suggests direct correspondence with a force representation, and we utilize such ideas below.

### Relating forces to the Hamiltonian

The force is related to the Hamiltonian (energy) by

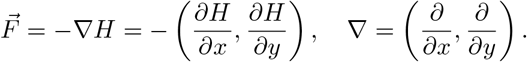

For the Hamiltonian given in the main text by (??), it has been noted by e.g., [5] that the same force can also be expressed in the form

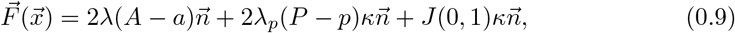

where, 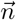 is the inward unit normal vector at the cell boundary and *κ* is the local curvature.

### Approximating forces at points along cell boundaries

For *h* = Δ*x* = Δ*y* the grid size and 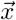 a point of a cell boundary, let *σ* be the cell configuration. Then a single “spin flip” at 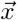 produces a small change in the Hamiltonian. This idea allows us to approximate *F_x_* = –*∂H*/∂*x* ≈ –Δ_*x*_*H*/Δ*x*, and similarly for *F_y_*.

CPM spin-flips are variations of the cell configuration, *σ*. We define *d_x_σ*, *d_y_σ* as spin-flips in the *x* and *y* directions that displace the cell boundary to the left or right relative to the given lattice site.

The centered difference approximation to the first partial derivative results in

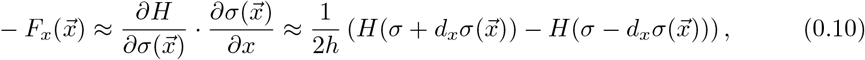

and similarly for the component –*F_y_*. Spin flips along any flat edge do not change to the configuration. Hence the direction of the force at such points would always be normal to the flat edge.

Using the fact that *H*(*σ* + *dσ*) = *H*(*σ*) + *dH*(*σ* → *σ* + *dσ*), we can rewrite the above as

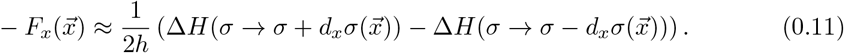

There are some special cases. If a site 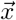 is positioned directly between two boundary points, as shown in Figure 11, then there are four possible spin-flips that affect the configuration at 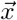: shifting the left-most cell edge out/in, or shifting the right-most edge out/in. These lead, respectively to the two approximations

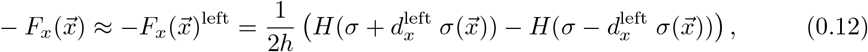

or

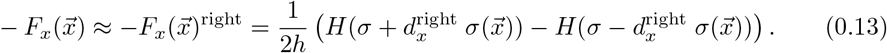

To avoid a bias in a particular direction, we resolve this by taking the average of Eqs. 0.12 and 0.13, so that

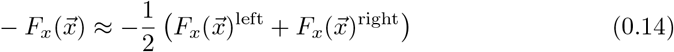

This approach is illustrated in Figure 11. Similar computations apply to *F_y_*, as before.

### Reducing the grid effects in perimeter calculations

Pixellation introduces artifacts in the perimeter of a cell. Approximating the cell perimeter as the sum of lattice edges (or number of lattice sites along the edge) is quite poor [2], introducing a large grid effect. We adopt the correction by [2] with the following neighborhood calculation:

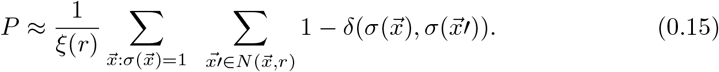

Here 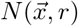 is the collection of neighboring sites of 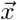 within a range *r* and ξ(*r*) is a scaling factor. (See Supplementary Figure 10.) This summation counts the number of neighboring sites of 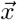 that are outside of the cell, i.e., how much 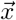 contributes to the perimeter. (Note that 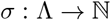 is the cell index or “spin value”, so that the delta function is nonzero only if 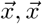 are both sites inside the same cell.)

A Moore neighborhood is often chosen (Figure 10) to compute the perimeter term (or adhesive energy term) in the Hamiltonian. For a neighborhoods with radius *r*,

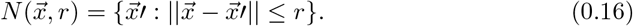

The larger the radius, the better the approximation of the perimeter (provided that the radius is not too large relative to cell size) [2]. Finally, the summation is normalized by the scaled factor ξ that corrects for the neighborhood radius [2]. We typically use a radius of 3 pixels for the neighborhood.

### Smoothing the forces

We refine the direction of the forces on the cell edge as follows. First we define 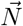 to be a weighted average of forces within a neighborhood as

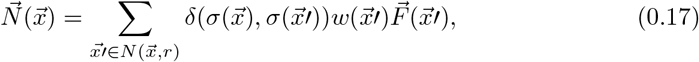

where

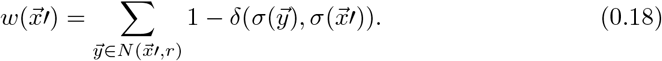

These weights count the number of neighborhood sites of 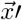 outside of the cell, i.e. the contribution of 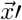 to the perimeter. We scale the vector 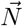 to obtain a unit vector 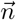, and then define the refined force as

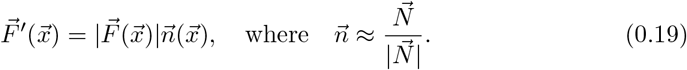

Here 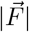 is the magnitude of the force given by our finite difference approximation. In Eq. 0.18, we use a neighborhood radius of *r* = 3, as recommended by [2].

### Optimal neighborhood size for smoothing

We numerically investigated the relationship between the smoothing neighborhood radius *r*, used in Eq. 0.17, and the accuracy of the smoothed force vector.

To do so, we took an elliptical shape, as in Supplementary Figure 12, for which the boundary normal vector is known.

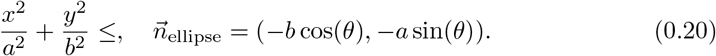

We first created a pixelated ellipse and displayed the approximated normal vectors for *r* = 3 in blue compared to the actual normal vectors of (0.20) (green). We next compared results of the smoothing algorithm of (0.17) with various values of the radius *r*. In each case, we compute the *L*^2^ norm (sum of squared errors, SSE),

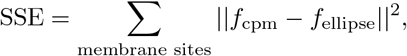

between the approximate (smoothed) normal direction and 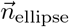. Supplementary Figure 12A shows the SSE for an ellipse with axes *a* =10 and *b* = 20 as a function of the radius *r*. We find that the optimal radius is *r* = 14. Panel D shows the smoothed normal vectors for *r* = 14 (blue) compared to actual normal vectors (green), showing that, indeed, there is improvement over a smoothing radius of *r* = 3.

We asked how the ellipse aspect ratios affect this conclusion. To test this, we varied the axes *a* and *b* of the ellipse, each from 7.5 to 47.5 in steps of 2.5. For each of these ellipses we computed the optimal *r* as before. Results are shown in Supplementary Figure 12C, where the elliptical axis *a* is on the *x*-axis and various values of *b* are shown with different colors. The optimal smoothing radius *r* increases roughly linearly with the length of either elliptical axis. Importantly, the SSE increases dramatically as *r* becomes too large relative to the cell size (Panel D), implying that larger *r* values are to be avoided. If cell shape is irregular, with small structures to be resolved, then large *r* values are inappropriate. We adopted *r* = 3 as a compromise.

### Phenomenological force fields in the interior

The centroid of the cell shape is

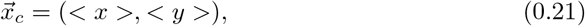

where < ·, · > denotes an average over all sites inside the cell. The goal is to define a force vector at every interior point 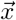 inside the cell. We do this by interpolating force vectors from the cell boundary to the centroid along straight line rays, assuming that net force at the centroid vanishes.

At each internal site 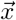 we identify the boundary site 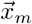 on such a ray to the centroid,

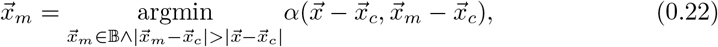

where 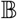 is the set of membrane sites and *α* denotes the angle between the given vectors. (See Supplementary Figure 13.) The force at 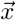 is assigned to be

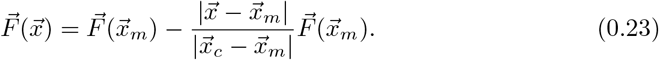

We used simple linear decay of force magnitude from boundary to centroid but othe assumptions such as quadratic, or exponential decrease are as easy to implement.

To smooth the vector field, we take an average over all boundary pixels that are neighbours to 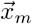:

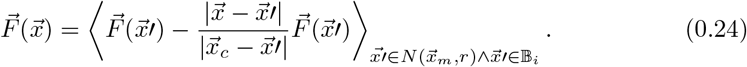

### Intracellular reaction-diffusion system and protrusive forces

For the internal signaling simulations, we used the wave-pinning reaction-diffusion model of [6]. Here *u*(*x*, *t*), *v*(*x*, *t*) are active and inactive forms of a signaling protein, e.g. Rho GTPase, satisfying the reaction-diffusion equations,

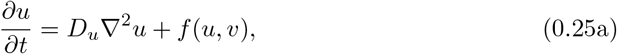

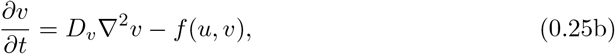

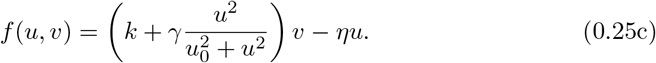

Parameters chosen were as follows for Figures 4 and 8: *D_u_* = 0.04, *D_v_* = 1, *η* = 5.2, *k* =1, *γ* = 30. Initial conditions were *u* = 0.04488, *v* = 0.4, with an elevated activity region with *u* = 4 along the left edge of the cell.

The reaction-diffusion equations are simulated in the irregular domain of the CPM cell, with 1000 iterations of the RD system per MCS. The spatial and time discretizations were *dx* = 0.03, *dt* = 0.00001.

We then assign a Hamiltonian difference *dH* to sites along the cell edge following the rule

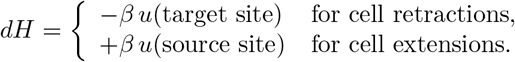

During each MCS, we update the chemical signaling field in the cell interior 1000 times. After an edge extension, we locally adjust the chemical distribution to avoid artifacts of numerical mass loss as follows: find all sites *x* within a range *r* of the source site *s*, let *T*(*u*) = Σ_*r*_*u* and *T*(*v*) = Σ_*r*_*v* be the total amounts of *u* and *v*, respectively, within that range; define a scaling factor

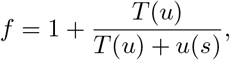

and similarly for *v*. Set the level of chemical activity in the new site (target site *t*) to *u*(*t*) = *u*(*s*) · *f*; at every one of the surrounding sites *x*, rescale *u*(*x*) · *f*. This ensures that the chemical level of the source site is copied into the target site, but the total level of signaling activity does not change.

After an edge retraction, we carry out a similar redistribution and scaling, but using the scaling factor

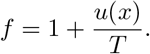

The level of active chemical *u* is then locally scaled by this factor.

We used different neighborhood balls for redistributing active versus inactive signaling levels, as the ranges of diffusion of these differ. Since *D_v_* = 25D*D_u_*, the range *r* is selected as *r_v_* = 25*r_u_*. Hence, the radii for redistribution were 3 for *u*, and 75 for *v*.

### Comparison to experimental data

The data for cell shape and traction force from [8] was provided to us on a triangular mesh. We first converted to the CPM square grid. Using the MatLab function “imresize”, the data was reduced to an 80 × 60 grid. Cell shapes were extracted by thresholding (Grey-scale values > 0.5 set to 1, < 0.5 set to 0, see Supplementary Fig. 17. Coordinates were scaled to –1 ≤ *x, y*, ≤ 1 in both data and CPM, and then images were superimpose. We identify the CPM coordinate closest to each data point and compared forces (CPM vs data) at these corresponding points.

The appropriate CPM parameters (λ, λ_*p*_, *J*(0,1), *A, P*) are not known a priori for the given cell types and conditions. These parameters are assigned as follows. First, we determined area and perimeter of the (scaled) data cells; these were found to be *a* = 1873, *p* = 1616.83 (round cell) and *a* = 949, *p* = 1271.94 (polarized cell). We then choose smaller target CPM area and perimeter for the given cell. Next, we select initial values for λ, λ_*p*_, *J*(0,1) such that all three corresponding terms in the Hamiltonian have a roughly similar contributions (λ = 0.5, λ_*p*_ = 1, *J*(0,1)=1000, *A*=500, *P* =1000.)

We first tested a smoothing radius of 3, as for our original method. Results are shown in Supplementary Figure 18. Because the direction of forces at protruding regions (circled) deviated strongly, we adopted a smoothing radius of 10. We rescaled the CPM parameters λ, λ_*p*_, *J*(0,1) by a constant scale factor *α* by minimizing the *L*^2^ norm between scaled CPM forces and data forces. This brings the force magnitudes to a common scale. We found that *α* ≈ 0.05, resulting in the parameters λ = 0.5 · *α* = 0.025, λ_*p*_ = 1 · *α* = 0.05, *J*(0,1) = 1000 · *α* = 50.

The above values of CPM parameters resulted in favorable comparisons between CPM forces and experimental data forces. However, we investigated whether a different CPM parameter set would lead to a better fit. To do so, we defined a range of parameter values: 0 < λ_*a*_, λ_*p*_ < 0.25, 0 < *J*(0,1) < 50, 0 < *A, P* < 2000, 0 < *r* < 50 (for membrane force smoothing). These ranges were binned into 10000 bins. We used a Latin Hypercube sampling, and sampled 100000 times. For each sample, we calculate the L2 norm between CPM and data forces. Overall, we found that different parameter sets gave very similar results. We used the first set in the Tables 1 and 2 for figures in the main text. In Supplementary Figure 22A we present box-plots of the angle between the model and experimental forces. Our method is slightly better at predicting the direction of traction forces for the circular cell, where traction force are more uniform.

**Table 1.** Top 5 parameters sets from the Latin hypercube sampling for the round cell, all giving very similar fits. The first set is used in the main text and the force fields for 2-5 are given in Supplementary Figure 19.

| rank | $\lambda$ | $\lambda_p$ | $J(0,1)$ | $A$ | $P$ | $r$ |
| --- | --- | --- | --- | --- | --- | --- |
| 1 | 0.0455 | 0.2122 | 45.15 | 1294.6 | 1802.4 | 31 |
| 2 | 0.0571 | 0.0827 | 12.755 | 1415.8 | 1908.2 | 31.3 |
| 3 | 0.0177 | 0.0925 | 37.98 | 404.6 | 1978.6 | 33.4 |
| 4 | 0.1123 | 0.1301 | 48.68 | 1637.8 | 1995 | 31.2 |
| 5 | 0.0213 | 0.1311 | 41.85 | 627 | 1979.6 | 31.6 |

**Table 2.** As in Table 1, but for the polarized cell. Multiple parameter sets give very similar fits with SSE around 1.3e6. First set is used in the main text and the force fields for 2-5 are given in Supplementary Figure 20.

| rank | $\lambda$ | $\lambda_p$ | $J(0,1)$ | $A$ | $P$ | $r$ |
| --- | --- | --- | --- | --- | --- | --- |
| 1 | 0.1716 | 0.0496 | 5.51 | 864.2 | 1423.8 | 12.2 |
| 2 | 0.0919 | 0.0608 | 24.43 | 797.8 | 1338 | 11.3 |
| 3 | 0.0355 | 0.1350 | 17.15 | 544.2 | 1329.2 | 12 |
| 4 | 0.0336 | 0.0529 | 15.89 | 527.8 | 1382.4 | 12.8 |
| 5 | 0.0573 | 0.1172 | 26.6 | 709.8 | 1397.4 | 12.7 |

In general, CPM parameters would, ideally, be optimized for a given cell type and conditions, and then used for predicting and validating other data not used in such optimization. Our data was limited, and so this optimization was beyond the scope of this initial series of tests.

### Multiple cells and forces on cell-cell interfaces

For multicellular aggregate, we decompose the total Hamiltonian into contributions *H^i^* made by each cell.

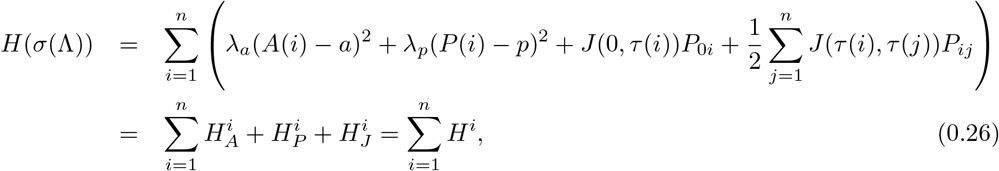

where *P*_0*i*_ is the length of the membrane of cell *i* that is in contact with the medium:

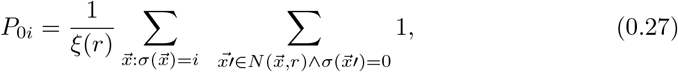

and *P_ij_* is the length of the interface between cell *i* and cell *j*:

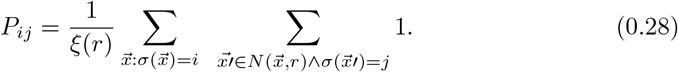

We use CPM spin-flips to calculate the force due to shifting the cell-cell interface (see Supplementary Fig. 23).

For a multicellular cluster at equilibrium, the force-balance principle states that traction forces must integrate to zero over the cell (or cell-cluster) footprint in 2D [18,29]. This can serve as an additional check on the predictions.

## Other Simulations of Multicellular aggregates

We start with a checkerboard type simulation as shown in Supplementary Figure 25. Here, the heterotypic adhesive forces are higher than the homotypic adhesive forces, so that cells of different type repel each other resulting in a checkerboard pattern. We provide zoom-ins of group of grey cells within the clusters. The forces between grey cells are high and repulsive. As time proceeds, this allows those cells to move away from each other and the green cells to push in between them. In the whole cluster, we observe a decrease in forces, indicating that the aggregate is going towards a force balance. The pattern stabilizes (see configuration at 5000 MCS in Supplementary Figure 26) but due to random fluctuations and pressure on the cells in the interior of the cluster, high forces appear and disappear at different spots in the clusters.

If the adhesive forces between green cells and the medium is very high, the grey cells will engulf the green cells. An example engulfment simulation is given in Supplementary Figure 27. Here the zoomed views track a region around the boundary of the cluster. Since green cells have high forces with the medium, they move into the clusters to avoid contact with the surrounding medium. The engulfment is not completed within the time frame shown here, but after 5000 MCS (Supplementary Figure 28) the engulfment is more or less complete.

